# Energy conversion mechanism revealed by ATP-free and ATP-dependent walking of myosin V motor

**DOI:** 10.1101/2025.10.11.681800

**Authors:** Noriyuki Kodera, Holger Flechsig, Takayuki Uchihashi, Toshio Ando

## Abstract

The mechanical processes of ATPase protein motors are coupled to their ATPase reactions. The free energy released at a given step(s) of the ATPase cycle is converted into mechanical work. Therefore, it has long been thought that ATP is the energy source for mechanical work. Here, we show that the double-headed myosin V (M5) motor can perform ATP-free walking. In the presence of ADP, M5 bound to F-actin by both heads makes a forward step when the trailing head is detached by an external force. This forward step harnesses the strain energy in the leading (L) head, provided from the strong binding energy between the L-head and actin. Consequently, strain generation reduces the binding affinity, which manifests as sporadic, brief L-head detachment. Strain generation and brief L-head detachment also occur during ATP-dependent walking to similar extents as during ATP-free walking. These results suggest that the strong binding energy is also converted into mechanical work during ATP-dependent walking.

---

ATP has been described as the energy source of all living cells. Indeed, ATPase motor proteins, including myosin, kinesin, dynein and F_1_, generate unidirectional force and movement by hydrolysing ATP, thereby performing a variety of cellular functions^1‒5^. The mechanical work is performed by a powerstroke^6,7^, which is defined as a rapid structural change that produces a large-scale motion comparable to the molecular size or as the formation of a highly strained conformation^8^. In the latter case, large-scale motion is generated by relaxation of the strain^8^.

This article examines the energy conversion mechanism in myosin V (M5), a double-headed processive motor protein that moves by hand-over-hand walking toward the plus end of F-actin with ∼36 nm advance per ATP hydrolysis cycle^9–13^. The term ‘head’ is defined as the motor domain plus the neck domain, while the term ‘lever-arm’ represents the neck domain plus the neck−motor domain junction (alias the converter hinge). Figure 1 illustrates a generally accepted chemomechanical cycle in M5 with a truncated tail (M5-HMM). M5-HMM will be referred to simply as M5. The cycle starts from the double-headed bound (DHB) state, where the trailing (T) head is nucleotide-free (NF) and the leading (L) head is bound to ADP. The neck domains of the T and L heads are oriented in forward and backward directions, respectively. However, the direction of the L-head neck region near the motor domain is often biased toward the forward direction, causing the entire neck to bend outward. This bending was observed by negative stain electron microscopy (nsEM) at low concentrations (0.5‒1.0 μM) of ATP^14^. At such low concentrations of ATP, ATP binding to the T-head is the primary rate-limiting step^15,16^, and therefore, M5 is predominantly in the initial state of the chemomechanical cycle (Fig. 1, top). ADP bound to the L-head is virtually unable to dissociate due to the strained structure of the L-head, which is the basis for processive hand-over-hand walking^13,17,18^.

**Fig. 1.**
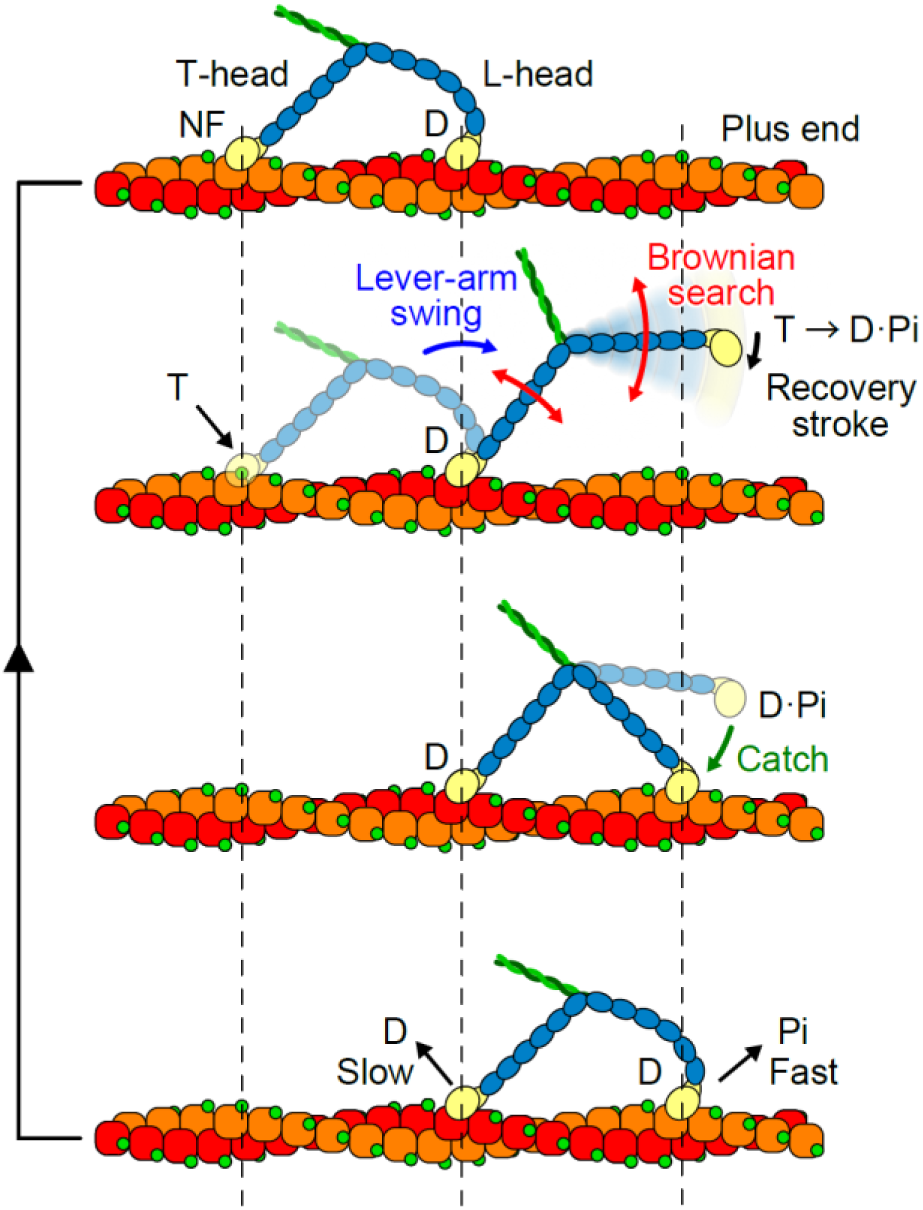
Generally accepted chemomechanical cycle in walking M5. NF, D, Pi, D·Pi, and T denote nucleotide-free, ADP, inorganic phosphate, ADP·Pi, and ATP, respectively. See the main text for details.

The chemomechanical cycle progresses as follows (Fig. 1). (1) Upon binding to ATP, the T-head detaches via a structurally revealed mechanism^19^, resulting in a forward swing of the L-head lever-arm through its strain relaxation. (2) The ATP bound to the detached head is rapidly hydrolysed into ADP and inorganic phosphate (Pi), resulting in a change in the angle between the motor and neck domains by 80‒90°^20–22^ via a structurally revealed mechanism^21,22^. This change in angle is known as the recovery stroke. It directs the actin-binding site of the motor domain to a forward actin site, thereby promoting head binding to actin in a backward-tilt orientation. (3) The detached head, undergoing Brownian motion around the forward-shifted neck−neck junction that is also undergoing Brownian motion, searches for a forward actin site. (4) Upon actual binding to an actin site weakly, the bound head becomes the new L-head in a primed pre-powestroke conformation. The structure of this transient primed state has recently been elucidated by time-resolved cryo-EM^23^. The step size, or the separation between the two motor domains bound to F-actin, which we refer to as the ‘motor domain (MD) separation’, is approximately 36 ± *n* × 5.5 nm, where *n* = −2, −1, 0, 1, 2, 3, 4 for phalloidin-free F-actin, and *n* = −1, 0, 1, 2 for phalloidin-stabilised F-actin^24^. The number of actin subunits contained within the MD-separation is *N*_m-m_ = 13 ± 2*n*. This discrete variation in actin sites available for head binding is the consequence of two factors: the nearly straight walking of M5 on F-actin with minimal azimuthal distortion in the DHB state^25^ and the temporal variation in the F-actin helical pitch due to cumulative angular disorder between adjacent actin subunits^24^. (5) Pi is released from the new L-head, which strengthens the L-head‒ actin binding. This strong binding leads to the generation of a powerstroke, which is associated with a bend-to-straight conformational change in the L-head^26,27^. However, the order of Pi release and force generation in any myosin class is currently under debate^28‒34^. Nevertheless, the crystal structure of myosin VI^29^ containing Pi in its putative escape route (termed ‘the backdoor tunnel’^35^) suggests that the force generation occurs when Pi has left from the active site yet remains at the tunnel exit^29,36^. We follow this structurally suggested scenario, since the weak binding between actin and M5 with ADP·Pi at the active site would hinder the generation of a strong force^36^. (6) ADP is released from the T head, resulting in an additional forward shift of the neck‒neck junction by 2‒6 nm^17,37^ and the restoration of the M5 molecule to its initial state (but with an advance by 36 ± *n* × 5.5 nm).

In the initial state (Fig. 1, top), the L-head is mainly strained through Pi release, which is reflected in its bending. This strain is used for the L-head swing after T-head detachment. However, nsEM^14^ and high-speed atomic force microscopy (HS-AFM)³^8^ have also observed L-head bending in the presence of ADP. HS-AFM has additionally detected occasional unfolding of the short coiled-coil tail of M5 in the DHB state, resulting in M5 monomerisation^38^. This unfolding was not caused by the AFM tip, as monomerised M5 was also observed in nsEM images of M5 purified by co-pelleting with F-actin^39^. Following tail unfolding, the monomerised L-head rotated forward in a manner similar to the swinging lever-arm motion observed in the presence of ATP^38^. Therefore, in the presence of ADP, the DHB M5 may potentially step forward when its T-head is mechanically detached, rather than by ATP binding. In the present study, we examined this possibility by using interactive HS-AFM (iHS-AFM), where the AFM tip was used for both molecular imaging and head detachment. Based on these and other observations, we investigated the energy conversion mechanism in ATP-dependent walking of M5.

## Results

### ATP-free walking of M5

In the present study, we used M5 from chick brain^40,41^. The assay system used for HS-AFM observation of M5 interacting with phalloidin-labelled F-actin immobilised on a lipid bilayer surface is identical to that employed previously^38^ (Fig. 2a and Methods). The lipid bilayer also contains 5% (w/w) of a positively charged lipid that weakly attracts M5 molecules to the bilayer surface. This attraction provides M5 molecules with a mechanical load against their movement along F-actin. This load reduces the translocation velocity by 27% at saturating concentrations of ATP^38^.

**Fig. 2.**
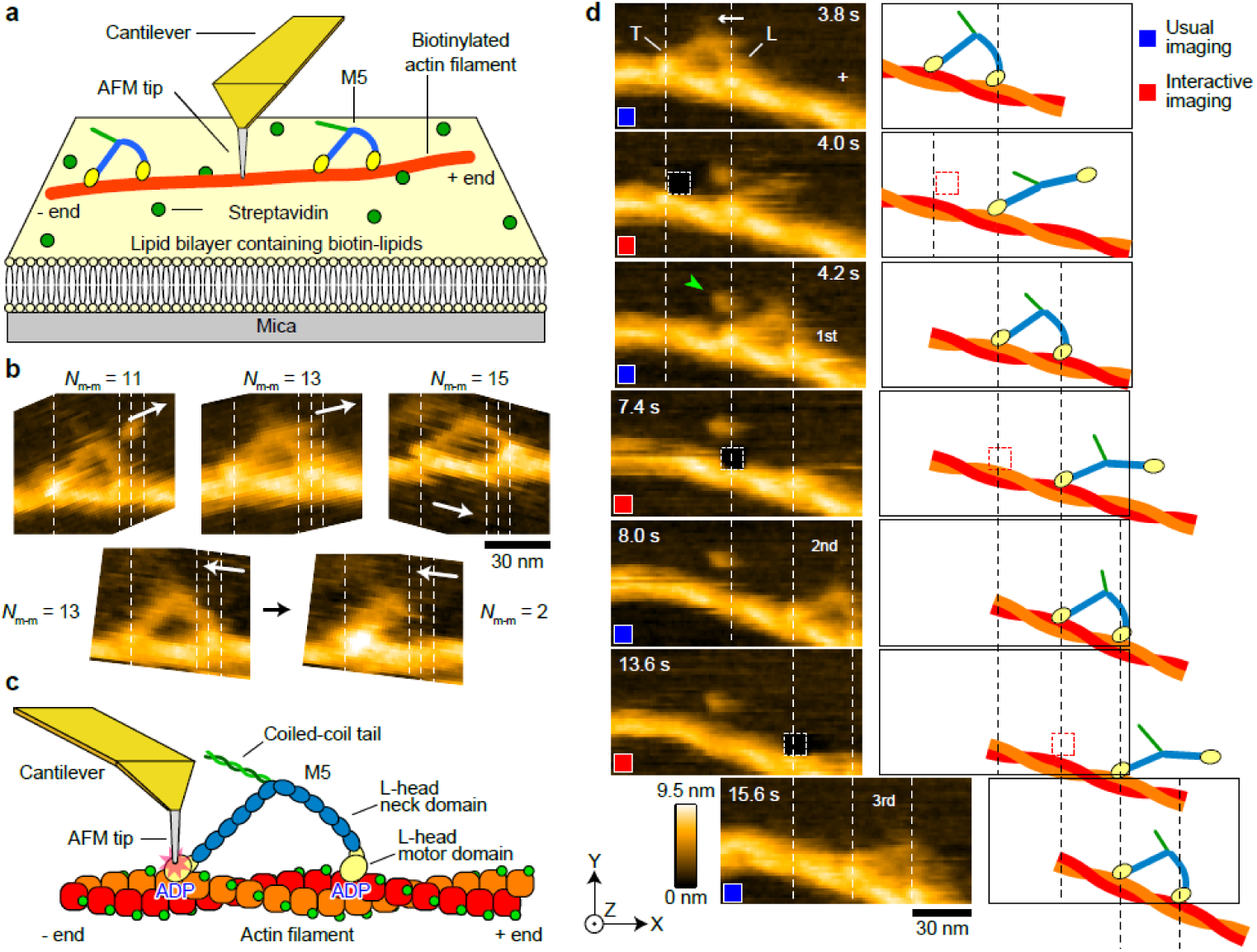
iHS-AFM imaging of ATP-free walking of M5. **a** Schematic of the assay system used for the imaging of ATP-free and ATP-dependent walking of M5 (see Methods). **b** Typical HS-AFM images showing actin-bound M5 with different MD-separations: top, in the presence of 1 mM ADP; bottom, in the presence of 1 µM ATP. The white arrows indicate the X-scanning directions during image acquisition. **c** Schematic showing the application of a strong tip force to an ADP-bound motor domain. **d** Typical successive HS-AFM images showing ATP-free walking of M5. Scan area, 130 × 65 nm^2^; number of pixels, 80 × 40 pixels; frame rate, 5 fps. A strong force was applied to the areas (7 × 7 pixels; dark rectangles) encircled with a white dashed line (4.0, 7.4 and 13.6 s). White arrow, coiled-coil tail of M5 tilted toward the minus end of F-actin; vertical dashed lines, the centres of mass of the motor domains; plus sign, the plus end of F-actin. The schematics on the right-hand side explain the AFM images, where the areas to which strong force was applied are marked with red dashed lines. The images were acquired while the tip was being scanned towards the +X and +Y directions during raster scanning.

In the presence of 1 mM ADP, M5 molecules were bound to F-actin by both heads with *N*_m-m_ = 11 (1.5%), 13 (90.7%), 15 (7.6%), and 17 (0%) actin subunits (typical AFM images are shown in Fig. 2b, top). M5 molecules with *N*_m-m_ = 2 also appeared by 0.12%, where the two heads were bound to adjacent actin subunits within the same single strand of double-stranded F-actin. Even in the presence of 0.1 and 1 μM ATP, this type of binding occurred to a similar extent as in the case of ADP. Its typical AFM image is shown in Fig. 2b, bottom, together with an AFM image of M5 with *N*_m-m_ = 13 that appeared before forming *N*_m-m_ = 2 at 1 μM ATP. For molecules with *N*_m-m_ = 2, we did not perform any additional experiment and analysis. The dynamic process leading to the DHB state could not be captured with HS-AFM, as this binding had already been completed by the time the HS-AFM system was ready for imaging after the addition of M5. Nonetheless, the observation of double-headed binding with *N*_m-m_ = 11, 13, and 15 suggests that the unbound head, which is destined to become the L-head, must have undergone a thermally induced angle change at the converter hinge prior to reaching the final state. Otherwise, the unbound head would have been unable to bind actin in the backward-tilt orientation. Although it is uncertain whether this angle change is similar to the structurally defined recovery stroke^21,22^, we refer to this thermally induced angle change as a thermal recovery stroke. Unexpectedly from the crystal structures of M5, previous nsEM observations of M5 alone revealed a NF head that was largely bent by ∼90° at the neck‒motor domain junction (resembling a post-recovery stroke conformation) as well as a straighter head at 100 μM ATP (resembling a post-powerstroke conformation)^42^. The appearance frequencies of these conformations were lower than those of opposite counterparts under the respective conditions^42^. Therefore, the neck‒motor domain junction is sufficiently flexible to permit a thermal recovery stroke to occur in the presence of ADP. However, upon binding to actin, the converter hinge angle that has been temporarily altered by a thermal recovery stroke rapidly redistributes itself around the most stable angle, which is restricted by the strong head‒actin binding. It can thus be inferred that L-head binding to actin following a thermal recovery stroke results in the L-head being subjected to strain. To ascertain whether ADP-bound M5 can advance by harnessing the strain energy acquired presumably in this way, we employed the iHS-AFM technique^43,44^, as explained below.

For the HS-AFM imaging of actin-bound M5 in the presence of 1 mM ADP, 0.1 µM ATP, or 1 µM ATP, the cantilever oscillation amplitude to be maintained during imaging (i.e., the set-point amplitude *A*_s_) was set to 0.9 × *A*_0_, where *A*_0_ (= 1.5 nm) is the free oscillation amplitude. Therefore, the mean oscillation energy loss per tip‒sample contact, *ΔE*_ts_, was estimated to be *ΔE*_ts_ = 3.1‒4.7 *k*_B_*T* (*k*_B_, Boltzmann constant; *T* = 298 in kelvin) from the mechanical properties of the short cantilevers used (see Methods). In iHS-AFM (Supplementary Fig. 1), which is used only in the presence of 1 mM ADP, the operator of the HS-AFM system specifies a pixel position within the successively displayed molecular images using a PC mouse pointer and then pushes a start key to initiate the interactive mode in the next frame of imaging. It should be noted that the T- and L-heads can be readily distinguished from one another. This is based on the curved L-head and also on a specific rule between the polarity of F-actin and the direction in which the heads bind to actin^45^ (Supplementary Fig. 2).

When the cantilever tip reaches a locus within a molecule corresponding to the specified pixel, *A*_s_ is rapidly reduced to a preset level *A*_p_. This results in the application of a controlled strong force to the specified locus by moving the sample stage vertically towards the oscillating tip. Immediately after this large force application, the system returns to the normal imaging mode. The HS-AFM system interprets this excessive vertical displacement as a very low height of the locus, such that the specified pixel appears darker in the image. In the experiments described below, *A*_p_ was set to 0.33 × *A*_0_ (= 0.5 nm), corresponding to Δ*E*_ts_ = 14.6‒22.0 *k*_B_*T*. Furthermore, to ensure 100% head detachment, the strong force was applied to closely positioned 7 × 7‒11 × 11-pixel points including the operator-specified point at their centre (for more details, see Supplementary Fig. 1). It should be noted that the oscillation energy, *ΔE*_ts_ = 3.1‒4.7 *k*_B_*T* or 14.6‒22.0 *k*_B_*T*, is transferred to the sample at each tip‒sample contact. However, this transferred energy dissipates rapidly (within much shorter than the oscillation period of ∼1 μs) into the surrounding water. Consequently, the transferred energy does not accumulate in the sample, as demonstrated by the HS-AFM imaging of the stator ring of F_1_-ATPase at 12.5 fps for 40 s using a cantilever with a resonant frequency of 1.2 MHz in water, where repeated conformational changes were continuously observed without any loss of activity^46^. In these conditions, a single stator ring was iteratively tapped with the tip tens of millions of times.

The application of a strong force to a motor domain (Fig. 2c) could detach the motor domain from actin with nearly 100% probability. The event of no detachment only rarely occurred but tended to occur when the motor domain shifted its position due to a change of the F-actin helical pitch that occurred by chance just prior to the initiation of the interactive mode. Since the vertically applied strong force pushes the motor domain toward its bound actin subunit, head detachment is very likely to be caused by brief structural deformation of the motor domain. When a strong force was applied to the L-head motor domain, the molecule appeared to remain in the same position despite brief detachment of the L-head. This detachment was confirmed by the brief disappearance of the motor domain from the image (Supplementary Fig. 3). In contrast, when the T-head motor domain was detached by the strong force, the molecule stepped forward with a high probability (92.5%). When this operation was repeated, the molecule almost always advanced in each interactive-mode operation, resulting in a long-range movement (Fig. 2d; Supplementary Video 1). It is noteworthy that the stepping direction was independent of the scanning direction (Supplementary Fig.4; Supplementary Video 2). Therefore, the observed forward stepping is not attributed to tip dragging. Furthermore, M5 retained its ATP-dependent motor activity after repeated application of a strong force, as evidenced by processive walking of such molecules after the addition of ATP by the UV photolysis of caged-ATP (Supplementary Fig. 5; Supplementary Video 3).

### Comparison of strain energies in different nucleotide conditions and MD-separations

The observed ATP-free walking indicates that the forward step of M5 uses the strain energy in the L-head provided just by the double-headed binding to F-actin. A crucial question is whether the mechanism of energy usage in ATP-dependent walking is identical to that in ATP-free walking. It remains unclear whether the strained L-head in the DHB state formed via the ADP·Pi-induced recovery stroke and subsequent Pi release is identical (or similar) to that formed via the thermal recovery stroke alone. To address this question, a quantitative comparison of the strain energies produced in the three nucleotide conditions (1 mM ADP, 0.1 µM ATP, or 1 µM ATP) was made. It is important to note that the chemical state of the T-head depends on the nucleotide conditions. In the presence of 0.1 and 1 µM ATP, the actin-bound T-head is predominantly NF, judging from the ADP dissociation rate constant of 7.2 s^‒1^ at the T-head under the experimental conditions^38^, whereas the L-head is bound to ADP in all three conditions.

In the DHB state, M5 is primarily subjected to strain at the converter hinges of the T and L heads, as well as at the neck domain of the L-head. To estimate the strain energy at each site, the distribution of the neck orientation angle (*φ*), relative to the F-actin orientation, of single-headed M5 bound to F-actin was first measured for both ADP-bound and NF states^38^ (Fig. 3a). The angular spring constants of the converter hinge *K*(*φ*), as estimated from the respective angle distributions *P*(*φ*), are dependent on the angle. For instance, at 1 mM ADP, the spring constant *K*_ADP_ is 23.0 *k*_B_*T*·rad^‒2^ at the equilibrium angle *φ*_0_ = 28.5°, decreasing with increasing *φ* (Fig. 3b, solid red line). In the NF state, the angular spring is stiffer. For instance, the spring constant *K*_NF_ is 40.0 *k*_B_*T*·rad^−2^ at *φ*_0_ = 28.5° and 37.9 *k*_B_*T*·rad^−2^ at the equilibrium angle in the NF state, *φ*_0_ = 32.9° (Fig. 3b, dashed red line). The potential energies of the converter hinge in the ADP-bound and NF states, *U*_D_(*φ*) and *U*_NF_(*φ*), are shown in Fig. 3b with the black solid and black dashed lines, respectively. The corresponding forces *F*(*φ*) generated, in parallel to F-actin, at the neck free end in the ADP-bound and NF states are shown in Fig. 3b with the blue solid and blue dashed lines, respectively. *F*(*φ*) in the ADP-bound state approaches a plateau (approximately 2.2 pN) at *φ* > 100°, which is consistent with the reported stall force for M5, 1.9–3.0 pN^47–49^. The derivation of these mechanical quantities is described in Methods.

**Fig. 3.**
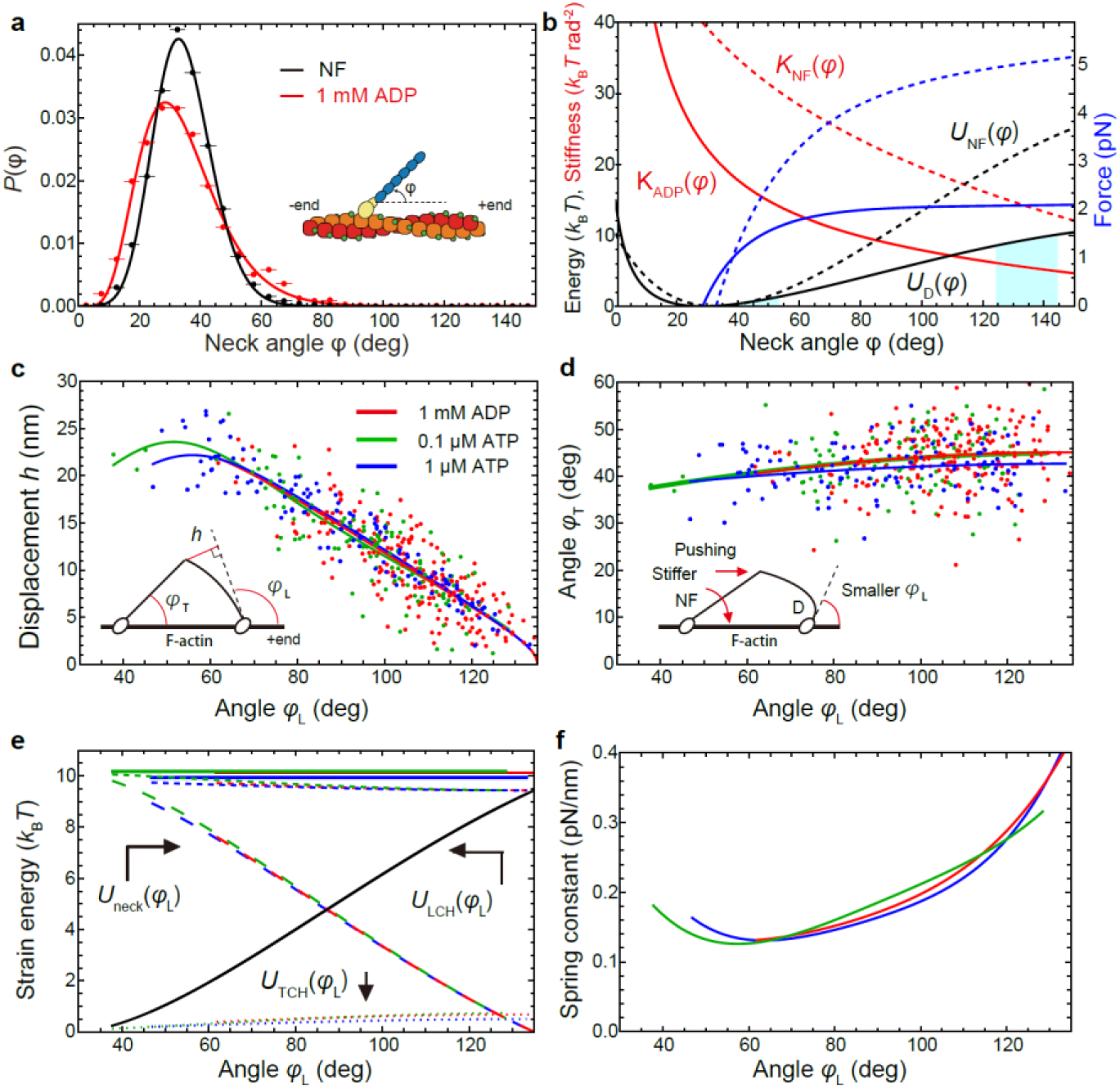
Mechanical properties of converter hinges and L-head neck of M5 in three nucleotide conditions. **a** Distributions of neck angle of actin-bound single-headed M5 in the NF and 1 mM ADP conditions. The horizontal bars indicate bin width 5°. **b** Angle-dependence of stiffness (red) and energy (black) of converter hinge, and the magnitude of force in the direction of F-actin generated at the free end of the neck (blue). The solid and broken lines are those for 1 mM ADP and NF conditions, respectively. **c−f** Mechanical properties of M5 with MD-separation of *N*_m-m_ = 13. The lines are drawn within the observed angle ranges of *φ*_L_. The colour code shown in (**c**) is common in (**c−f**), except for the black line in (**e**). In (**c**), lever-arm displacement *h* depending on the angle *φ*_L_ is shown. The solid lines were obtained by fitting the plotted data to fourth polynomial functions. The inset shows three geometrical parameters (*φ*_L_, φ_T_, and *h*) measured. In(**d**), dependence of *φ*_T_ on *φ*_L_ is shown. The solid lines were obtained by fitting the plotted data to second polynomial functions. The inset shows the stiffer NF T-head pushing the neck−neck junction. In (**e**), strain energies as a function of *φ*_L_ are shown. *U*_TCH_ and *U*_LCH_ are strain energies of the converter hinges at the T and L heads, respectively. *U*_neck_ is the bending energy of the L-head neck. The top horizontal solid lines show the total strain energy (*U*_TCH_ + *U*_LCH_ + *U*_neck_), and the dashed lines slightly below the horizontal solid lines show *U*_LCH_ + *U*_neck_. In (**f**), spring constants of L-head neck bending, *k*_neck_, are shown.

Subsequently, the geometric parameters of M5 in the DHB state were measured: the tangential angle of the L-head neck region emerging from the motor domain, *φ*_L_, the angle of the T-head neck as a function of *φ*_L_, *φ*_T_(*φ*_L_), and the deflection of the L-head neck end (at the neck−neck joint) as a function of *φ*_L_, *h*(*φ*_L_) (see Fig. 3c, inset). The experimental data of *h* = *h*(*φ*_L_) and *φ*_T_ = *φ*_T_(*φ*_L_) were fitted to the fourth and second polynomial functions, respectively (solid lines in Fig. 3c,d for *N*_m-m_ = 13 actin subunits and in Supplementary Fig. 6a-d for *N*_m-m_ = 11 and 15 actin subunits). Note that each solid line is drawn in a range of observed angles of *φ*_L_. The maximum angle of *φ*_L_ (denoted with *φ*_L_^max^), where the L-head neck deflection was extrapolated to zero, was approximately 127° for *N*_m-m_ = 11, 135° for *N*_m-m_ = 13, and 145° for *N*_m-m_ = 15, irrespective of the nucleotide conditions. For each MD-separation, the fitted curves for *h* = *h*(*φ*_L_) are nearly identical across the three nucleotide conditions (the three solid lines in Fig. 3c for *N*_m-m_ = 13 and those in Supplementary Fig. 6a,b for *N*_m-m_ = 11 and 15). This is also the case for *φ*_T_ = *φ*_T_(*φ*_L_) (the three solid lines in Fig. 3d for *N*_m-m_ = 13 and those in Supplementary Fig. 6c,d for *N*_m-m_ = 11 and 15). The only distinct difference between the nucleotide conditions is that the minimum value of *φ*_L_ that the L-head can take is smaller (approximately by 20−30°) in the presence of ATP than in the presence of ADP (see the left end of each fitted solid lines in Fig. 3c,d and Supplementary Fig. 6a−d). Since *φ*_T_ is always larger than the equilibrium angles *φ*_0_ of single-headed M5 (28.5° for the ADP-bound head, and 32.9° for the NF-head) irrespective of the nucleotide conditions, the stiffer NF T-head at 0.1 and 1 µM ATP pushes the ADP-bound L-head at the neck-neck junction more strongly than the less stiff ADP-bound T-head at 1 mM ADP (Fig. 3d, inset), resulting in the smaller minimum value of *φ*_L_ in the presence of ATP. This is how the 2-6 nm forward shift of the neck−neck (or neck−tail) junction occurs when ADP detaches from the T-head^17,37^.

The DHB state remains stationary under all nucleotide conditions, except for thermal fluctuations. Consequently, all forces acting on a M5 molecule in the DHB state, including the constraining forces, are balanced. As such, the sum of the strain energies at the three sites is conserved (from the principle of virtual work in mechanics; see Methods). When the strain energies at the converter hinges of the T and L heads and at the neck domain of the L-head are denoted as *U*_TCH_, *U*_LCH_, and *U*_neck_, respectively, the sum *U*_TCH_[*φ*_T_(*φ*_L_)] + *U*_LCH_(*φ*_L_) + *U*_neck_[*h*(*φ*_L_)] is constant (≡ *E*_total_) over possible values of *φ*_L_, under a given condition. The value of *U*_TCH_[*φ*_T_(*φ*_L_)] can be estimated from *U*_D_(*φ*) or *U*_NF_(*φ*) at *φ* = *φ*_L_ (solid or dashed black lines in Fig. 3b) depending on the nucleotide condition, while the value of *U*_LCH_(*φ*_L_) can be estimated from *U*_D_(*φ*) at *φ* = *φ*_L_ (solid black line in Fig. 3b). The resulting *U*_TCH_[*φ*_T_(*φ*_L_)] and *U*_LCH_(*φ*_L_) for *N*_m-m_ = 13 are shown in Fig. 3e with the dotted lines and the black solid line (common to all nucleotide conditions, except for the angle ranges), respectively, and those for *N*_m-m_ = 11 and 15 are shown in Supplementary Fig. 6e,f. The value of *E*_total_ can be obtained as *E*_total_ = *U*_TCH_[*φ*_T_(*φ*_L_^max^)] + *U*_LCH_(*φ*_L_^max^). The results are shown in Fig. 3e with solid horizontal lines for *N*_m-m_ = 13, while those for *N*_m-m_ = 11 and 15 are shown in Supplementary Fig. 6e,f. As can be clearly seen by the solid horizontal lines in these figures, the total strain energy is very similar (around ∼10 *k*_B_*T*) across the three nucleotide conditions. It is also similar across the different MD-separations. This similarity across the different MD-separations is consistent with the report that there is no significant difference in the kinetics of ATP-dependent walking regardless of the stride length^24^. The strain energy used for the mechanical work performed by the L-head swing after T-head detachment is not the total strain energy *E*_total_ but *U*_LCH_(*φ*_L_) + *U*_neck_[*h*(*φ*L)] (dashed lines located just below the solid horizontal lines in Fig. 3e and Supplementary Fig. 6e,f). However, the contribution of *U*_TCH_[*φ*_T_(*φ*_L_)] to *E*_total_ is small and almost identical across the three nucleotide conditions, although the maximum value of *U*_TCH_[*φ*_T_(*φ*_L_)] differs across the different MD-separations (0.5 *k*_B_*T* for *N*_m-m_ = 15, 0.7 *k*_B_*T* for *N*_m-m_ = 13, and 1.5 *k*_B_*T* for *N*_m-m_ = 11). Thus, the strain energy to be used for the forward step is very similar between ATP-free walking and ATP-dependent walking, despite the different pathways to the strained state as described above.

The value of *U*_neck_[*h*(*φ*_L_)] can be obtained as *U*_neck_[*h*(*φ*_L_)] = *E*_total_ − *U*_TCH_[*φ*_T_(*φ*_L_)] − *U*_LCH_(*φ*_L_) (long dashed lines in Fig. 3e and Supplementary 6e,f). The spring constant of the L-head neck, *k*_neck_, estimated from *U*_neck_[*h*(*φ*_L_)] depends on *φ*_L_ and the MD-separation (Fig. 3f and Supplementary Fig. 6g,h). These dependencies indicate that larger neck bending (i.e., smaller *φ*_L_) results in smaller *k*_neck_. The value of *k*_neck_ varies around 0.18 pN/nm, a value previously estimated using optical tweezers^17^. Note that the estimated values of the spring constant in the large *φ*_L_ regions are not reliable because the values were obtained by dividing the values of 2*U*_neck_ by small values of *h*^2^.

### Foot stomping

If a M5 molecule is bound to actin solely by the T-head, the molecule will not undergo strain except for that induced by thermal effects. Therefore, the total strain energy in M5 in the DHB state at 1 mM ADP originates from the strong binding energy between the L-head and actin. Interestingly, it was observed at 1 mM ADP that the L-head occasionally detached from actin and then quickly reattached to F-actin at the same position or its vicinity (Supplementary Fig. 7a; Supplementary Video 4). This behaviour, which occurs predominantly at the L-head, was previously observed by HS-AFM also in the presence of 0.1 and 1 µM ATP^38^. This behaviour is referred to as ‘foot stomp’. In our previous study, positional shifts of the motor domains approximately by ±5.5 nm without detachment (Supplementary Fig. 7b,c; Supplementary Vide 5) were misclassified also as foot stomp. The positional shifts without detachment are now referred to as ‘foot sliding’ and are understood to result from changes in the helical pitch of F-actin. The foot sliding was observed to occur at either head (L-head, Supplementary Fig. 7b; T-head, Supplementary Fig. 7c), resulting in wider or shorter MD-separations. Foot stomping is not due to tip force, as similar behaviour has been observed during ATP-dependent M5 walking using interferometric scattering (iSCAT) microscopy^50^ and polarised fluorescence microscopy^51^. The frequency of foot stomping observed with iSCAT was ∼3% of steps taken at 1 µM ATP, ∼0.6% of steps taken at 10 µM ATP, and none at saturating ATP^50^. The lower frequency at higher ATP concentrations is likely attributable to the T-head being detached by its binding to ATP during L-head foot stomping, resulting in complete dissociation of the molecule. Such foot stomp events would have been excluded from the count. This complete dissociation is likely to be one of the mechanisms by which processive walking of M5 is terminated. Since single-headed M5‒ADP has a high affinity for actin (*K*_d_ = 7.6 × 10^−9^ M, corresponding to a binding energy of –18.7 *k*_B_*T*)^16^, foot stomping at the L-head is most likely caused by a reduction in its binding energy equivalent to the total strain energy of ∼10 *k*_B_*T*. Therefore, the affinity of the L-head reduces to approximately *K*_d_ = Exp[(–18.7 + 10)] ≈ 1.7 × 10^−4^ M in the DHB state under the three nucleotide conditions, which must manifest as foot stomping. We verified this issue by estimating the foot stomp rate (*R*_stomp_) by counting the number of foot stomp events (*N*_stomp_) out of the total number of imaged frames (*N*_total_), as *R*_stomp_ = *N*_stomp_/*N*_total_ × imaging rate, where *N*_stomp_ observed at different MD-separations were summed. This summation is because the total strain energy is similar between different MD-separations. As shown in Table 1, *R*_stomp_ was similar across the three nucleotide conditions (0.069 s^−1^, 0.064 s^−1^, and 0.109 s^−1^ at 1 mM ADP, 0.1 μM ATP, and 1 μM ATP, respectively). Our results of *R*_stomp_ are similar to those estimated from the frequencies of foot stomping observed with iSCAT microscopyy^50^, although some foot stomp events are likely to be missed at 10 µM ATP due to complete dissociation of M5: 0.05 s^−1^ and 0.04 s^−1^ at 1 µM and 10 µM ATP, respectively. Here, the second-order ATP binding rate constant (2.0 μM^−1^s^−1^) and the ADP dissociation rate constant (10.8 s^−1^)^38^ at the T-head were used to estimate *R*_stomp_ from the foot stomp frequencies observed by iSCAT microscopy^50^. Thus, the generation of molecular strain reduces the affinity of the L-head for actin, resulting in foot stomping, while the generated strain energy *U*_LCH_(*φ*_L_) + *U*_neck_[*h*(*φ*_L_)] is converted into mechanical work performed by the actin-bound L-head during both ATP-free and ATP-dependent walking.

**Table 1.** Foot stomp events observed under three nucleotide conditions.

| Nucleotide conditions | Total number of images | Number of foot stomp events | Imaging rate (fps) | Foot stomp rate ($s^{-1}$) |
| --- | --- | --- | --- | --- |
| 1 mM ADP | 25,790 | 588 | 3.0 | 0.068 |
| 0.1 μM ATP | 7,441 | 71 | 6.67 | 0.064 |
| 1 μM ATP | 5,709 | 93 | 6.67 | 0.109 |

Although very rare, M5 exhibited 30 instances of a backward step following spontaneous L-head detachment, among the 25,790 images captured in the presence of 1 mM ADP. To counteract this backward step, M5 displayed 28 forward steps following spontaneous T-head detachment in the presence of 1 mM ADP. The similar frequencies of the forward and backward steps caused by spontaneous head detachment are therefore consistent with the principle of detailed balance under thermal equilibrium.

### Energetics of ATP-free and ATP-dependent walking

This section uses an energy landscape to quantitatively explain the relatively low success/failure ratio in ATP-free walking compared to ATP-dependent walking. This landscape leads to the landing of the detached head on F-actin through molecular processes occurring at the detached and attached heads. The results show the ADP·Pi-induced recovery stroke greatly contributes to the high degree of successful forward stepping in ATP-dependent walking, despite a small free energy being expended for the recovery stroke.

In ATP-free walking, the T-head, detached from actin by the strong tip force, reaches the next forward binding site through the following stages: (i) thermal recovery stroke by the detached head, (ii) forward swing of the actin-bound L-head, and (iii) Brownian search of the detached head for the next binding site^47,50,52,53.^. The Brownian search is performed by rotational diffusion of the bound head around the equilibrium angle relative to F-actin and by rotational diffusion of the detached head around the neck–neck junction (see Fig. 4a,b). However, the energy spent in the latter Brownian rotation is negligible^54,55^ because the neck–neck joint is highly flexible as indicated by the wide distribution of the angle made by the two necks at the joint^56^. Thus, the following equation holds for the success/failure ratio (= 0.925/0.075) of ATP-free forward stepping:

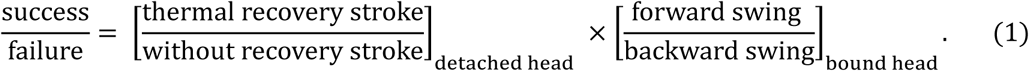

Here, the denominators and the numerators on the right-hand side represent the probabilities of respective molecular actions. From Eq.(1), the overall energy barrier difference between the forward step and the backward return after T-head detachment (Δ*E*_0_) is equal to the sum of the energy barrier differences corresponding to the first term (Δ*E*_1_) and the second term (Δ *E*_2_) on the right side of Eq.(1), i.e.,

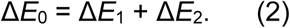

In the failure events, the detached motor domain returned to the original actin site or its vicinity. Despite the low probability of backward return, the distribution of MD-separations formed immediately after backward return was similar to that of forward step (pink and red bars in Supplementary Fig. 8). This similarity indicates that the distribution of different MD separations is predominantly determined by the probabilities that the half pitch of F-actin takes approximately 36 ± *n* × 5.5 nm. Even when the resulting MD separation differs between the two DHB states formed immediately after the success and failure steps, the two states are energetically similar, as the total strain energy (*E*_total_) is only slightly sensitive to the MD separation (Fig. 3e and Supplementary Fig. 6e,f). Consequently, the energy landscapes for the forward step and the backward return can be depicted as shown in Fig. 4c, where G_f_^ǂ^ and G_b_^ǂ^ denote the energy barrier heights for the forward step and the backward return, respectively. Their difference is given by G_f_^ǂ^ − G_b_^ǂ^ = Δ*E*_0_.

**Fig. 4.**
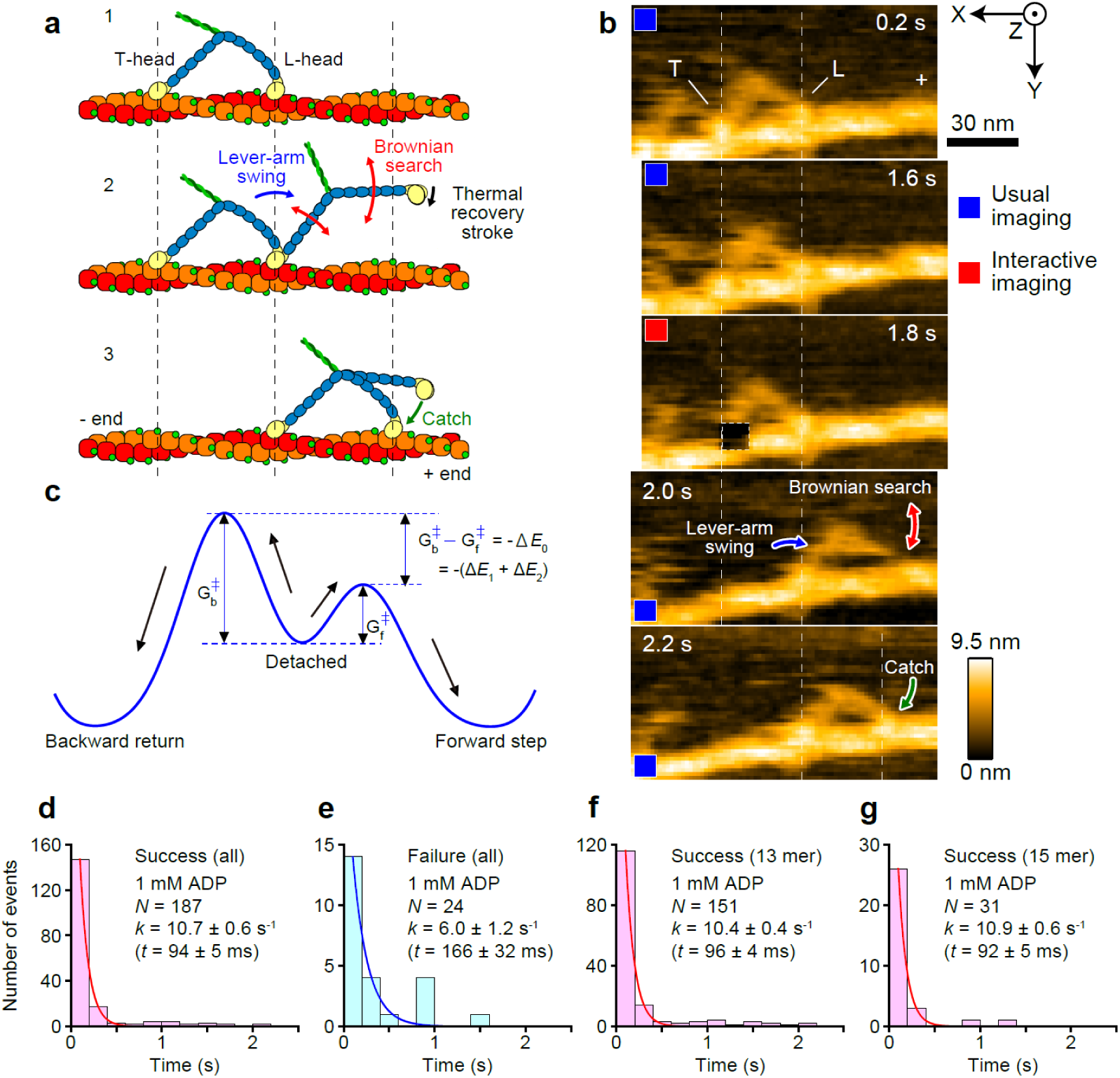
Transition events in ATP-free walking of M5. **a** Schematic showing transition events occurring between the T-head detachment and its final binding to a front actin site. **b** Energy landscape leading the detached head to forward step or backward return. Both the activation energies (G_f_^ǂ^ and G_b_^ǂ^) are assumed to contain an identical unspecifiable component in addition to *E*_1_ (thermal recovery stroke energy only for the forward step) and *E*_2_ (energy for lever-arm swing). **c** Typical successive HS-AFM images showing transition events. Scan area, 130 × 65 nm^2^; number of pixels, 80 × 40; frame rate, 5 fps. A strong force was applied to the area (7 × 7 pixels; dark rectangle) encircled with a white dashed line (1.8 s). The images were acquired while the tip was being scanned towards the +X and +Y directions during raster scanning. **d-g** Histograms of time duration between T-head detachment by tip-force and its landing on a front actin site (**d,f,g**) or on an original actin site or its vicinity (**e**). The red and blue lines indicate a single exponential function fitted to the respective histograms. Note that the time resolution is limited to the image acquisition time (200 ms). In (**d,e**), all data are combined without distinguishing the MD-separations after landing. In (**f**), only data with *N*_m-m_ = 13 are combined. In (**g**), only data with *N*_m-m_ = 15 are combined.

The overall energy barrier difference Δ*E*_0_ is given by Δ*E*_0_ = –*k*_B_*T* log_e_(success/failure) = –2.5 *k*_B_*T* (see Fig. 4c). Next, we estimate the energy barrier difference Δ*E*_1_ between the presence and absence of a thermal recovery stroke by the detached head; see the first term on the right-hand side of Eq.(1). In ATP-dependent walking, the recovery stroke in the detached head is caused by the ATP hydrolysis into ADP·Pi in the detached head (Fig. 1)^20–22^ in a structurally well-defined manner^21,22,57^. The recovery stroke angle Δ*φ*_rs_ was previously measured to be 85° ± 19°^20^ using a bright-field optical microscope in combination with a UV-flash photolysis system for the release of ATP from caged-ATP. In this measurement, the neck of monomeric M5 was immobilised on a top surface of a bead, while the motion of a double bead specifically attached to the motor domain was observed in real time. The angular spring constant of the neck‒motor domain junction in the actin-unbound head was estimated to be *k*’ = 23 pN·nm·rad^−2^ = 5.59 *k*_B_*T*·rad^−2^ from the Gaussian width of the thermally driven angular fluctuations^20^, which was very similar between before and after a recovery stroke. The value of the angular spring constant is also similar to that of muscle myosins^58^. It is noteworthy that the spring constant *k*’ is significantly smaller than those of the single-headed M5 bound to actin in the ADP-bound and NF conditions, *K*_ADP_ and *K*_NF_, respectively (Fig. 3b). Thus, actin binding largely stiffens the converter hinge. Supposing that the value of Δ*φ*_rs_= 85° is the most adequate angle to achieve the MD-separation of 36 nm, Δ*φ*_rs_ ranges from 77° to 95° for covering the MD-separation range of 36 ± 5.5 nm. The mean thermal recovery stroke energy is approximately estimated to be Δ*E*_1_ = 0.5 × *k*’ × 〈Δ*φ*_rs_^2^〉 = 6.3 *k*_B_*T*, where averaging is made over the angle range of 77°–95°. As a consequence of this large uphill process, the forward landing of the detached T-head is slow (approximately 10.7 s^−1^, corresponding to a characteristic time ≈ 94 ms, Fig. 4d), compared to that observed in ATP-dependent walking (160–180 s^−1^)^47,52^. This result of the landing rate was obtained without considering the different MD-separations formed immediately after landing. For the kinetic scheme of

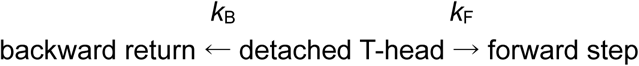

with rate constants *k*_B_ (for backward return) and *k*_F_ (for forward step), the landing rate *k*_landing_ should be the same between the two landing processes, i.e. *k*_landing_ = *k*_F_ + *k*_B_; from *k*_F_/*k*_B_ = 0.925/0.075, we obtain *k*_F_ = 9.9 s^−1^ and *k*_B_ = 0.80 s^−1^. However, the backward return landing was slower (about 6.0 s^−1^) than the forward landing, as estimated from the histogram of backward return times (Fig. 4e). This inconsistency is most likely due to the limited number of backward return events, as well as the limited time resolution. The relationship between the forward landing rate and the MD-separation (Fig. 4f,g) will be described later. From the values Δ*E*_0_ = –2.5 *k*_B_*T* and Δ*E*_1_ = 6.3 *k*_B_*T*, the energy barrier difference between the forward and backward swings of the actin-bound head can be indirectly estimated to be Δ*E*_2_ (= Δ*E*_0_ – Δ*E*_1_) = –8.8 *k*_B_*T*. However, the value of Δ*E*_2_ can be directly estimated, as described below. Just before forward landing of the detached head, the actin-bound head with ADP makes an angle of 45° ± δ relative to F-actin, while just before backward return landing, it makes an angle of 135° ± δ, where δ is approximately 10°, estimated from the differences of *φ*_L_^max^ between different MD-separations. Therefore, Δ*E*_2_ can be estimated as the mean potential energy difference, 〈*U*_D_(45°)〉 – 〈*U*_D_(135°)〉, where averaging is made over the angle regions, 45° – δ ≤ *φ* ≤ 45° + δ for 〈*U*_D_(45°)〉 and 135° – δ ≤ *φ* ≤ 135° + δ for 〈*U*_D_(135°)〉; see the light-blue regions under the solid black line in Fig. 3b. The values of 〈*U*_D_(45°)〉 and 〈*U*_D_(135°)〉 are 0.7 *k*_B_*T* and 9.4 *k*_B_*T*, respectively, which yields Δ*E*_2_ = –8.7 *k*_B_*T*. This result is in good agreement with the value indirectly estimated above. Thus, the energy landscape derived from Eq.(1) could quantitatively explain the observed success/failure ratio of the forward step in ATP-free walking.

In ATP-dependent walking, failure of the forward step is very rare. This is due to the first term on the right-hand side of Eq. (1) changing to the following form:

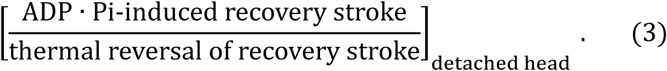

The second term on the right-hand side of Eq.(1) is common to both ATP-free and ATP-dependent walking. The energy barrier difference Δ*E*_1_ corresponding to the new form (3) becomes Δ*E*_1_ = – 2.3 *k*_B_*T* – (+ 6.3 *k*_B_*T*) = –8.6 *k*_B_*T*. Here, the –2.3 *k*_B_*T* is for the ADP·Pi-induced recovery stroke estimated from the ATPase kinetics^16^, while 6.3 *k*_B_*T* is the energy required to thermally reverse the ADP·Pi-induced recovery stroke (i.e., 0.5 × *k*’ × 〈Δ*φ*_rs_^2^〉). By including the second term shown in Eq.(1), the overall energy barrier difference between the forward step and the backward return becomes Δ*E*_0_ = –8.6 *k*_B_*T* – 8.7 *k*_B_*T* = –17.3 *k*_B_*T*. The success/failure ratio therefore becomes Exp(–17.3) ≈ 3 × 10^−8^. The ADP·Pi-induced recovery stroke largely contributes to this high success rate by elevating the energy barrier height for the backward return, despite the small amount of free energy expended (2.3 *k*_B_*T*).

### Mechanical model and simulations of ATP-free M5 translocation

We constructed a simplified mechanical model of the M5 dimer considering converter-hinge energies derived from HS-AFM data for the angular distribution *P*(*φ*) mentioned above (Methods, Supplementary Information). Brownian dynamics simulations of the model were employed to numerically investigate ATP-free M5 translocation cycles along F-actin (Methods, Supplementary Methods, Supplementary Fig. 9, and Supplementary Table 1). We investigated the following questions: (i) Is the external energy applied with the AFM tip under iHS-AFM scanning sufficient to induce mechanical detachment of the T-head? (ii) How long must the rebinding process for a step in an ATP-free cycle be?

As described above, the total strain energy is ∼10 *k*_B_*T* for M5‒ADP in the DHB state. To enable binding of an M5 head, the actin-binding energy should at least compensate for this total strain. Thus, the head‒actin binding energy is estimated to be at least –10 *k*_B_*T*. In the ATP-free case, the energy required for mechanically detaching the T-head has to be supplied by the AFM tip, which under interactive mode scanning provides sufficient energy (14.6‒22.0 *k*_B_*T* oscillation energy loss per tap). Importantly, the externally supplied energy is always larger than the amount that is temporarily borrowed from the environment during the thermal recovery stroke (Δ*E*_1_ = 6.3 *k*_B_*T* for Δ*φ*_rs_ = 85°), such that the energy balance for an entire forward step is consistent with the second law of thermodynamics (that is, the energy balance confirms that the steps proceed downhill).

Brownian dynamics simulations of the ATP-free M5 stepping revealed a forward landing time of 82 ms (rate, 12.2 s^−1^; *n* = 1852), while the backward return landing time was 110.4 ms (rate, 9.06 s^−1^; *n* = 148). When the loading effect of the substrate surface in the HS-AFM experiment is considered, these values become 112 ms (rate 8.9 s^−1^) and 151 ms (rate, 6.6 s^−1^) for the forward and backward return landings, respectively, which are roughly consistent with the HS-AFM data (94 ms for forward landing and 166 ms for backward landing). The step time distribution together with an exemplary time trace of the M5 for consecutive ATP-free stepping and corresponding conformational snapshots of the mechanical M5 model are shown in Supplementary Fig. 10 (also see Supplementary Video 6). Furthermore, the success/failure ratio for a forward step after mechanical T-head detachment obtained from simulations as 0.926/0.074 agrees remarkably well with HS-AFM observations of 0.925/0.075.

Although the simplified mechanical model of ATP-free M5 translocation does not resolve the detailed physical processes underlying an M5 step upon mechanical T-head detachment (and therefore does not allow us to infer their corresponding energetics), the numerical investigations confirm the HS-AFM observations that the process of rebinding and strain generation in the L-head does not require the execution of ATP-related conformational changes (i.e., the recovery stroke and the powerstroke) and corresponding chemical energy input.

### MD-separations after ATP-free and ATP-dependent forward steps

Although the MD-separation formed after forward stepping is not directly related to the main subject of this study (i.e., the energy conversion mechanism), two observed results and their interpretations are briefly described here. The first result is that two MD-separations of *N*_m-m_ = 13 and 15 showed similar landing rates in the ATP-free walking (10.4 s^−1^ for *N*_m-m_ =13 and 10.9 s^−1^ for *N*_m-m_ = 15; Fig. 4f,g). The number of events with *N*_m-m_ = 11 was too small to be analysed. The landing rate is influenced by the energy required for the thermal recovery stroke, 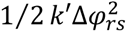, and the energy required for the forward orientation of the bound head *U*_D_(*φ*_L_), the sum of which is not significantly different between the two MD-separations: 6.87 *k*_B_*T* for *N*_m-m_ = 13 (Δ*φ*_rs_ = 85° and *φ*_L_ = 45°) and 7.83 *k*_B_*T* for *N*_m-m_ = 15 (Δ*φ*_rs_ = 95° and *φ*_L_ = 35°). This small difference, ∼1 *k*_B_*T*, is roughly consistent with the small difference in the landing rate, considering the insufficient time resolution of HS-AFM imaging, as well as the limited number of stepping events with *N*_m-m_ = 15.

The second result is the distribution of MD-separations taken immediately after forward stepping in ATP-free and ATP-dependent walking (Supplementary Fig. 8). The MD-separation is largely determined by the periodicity of F-actin that guides the positions of the L and T heads^36^. However, a discernible tendency is evident in the distributions across the three nucleotide conditions, albeit small differences. That is, the probability of taking a step with a MD-separation of *N*_m-m_ = 15 is greater in the presence of ATP than in the presence of ADP. Additionally, the probability of having *N*_m-m_ = 15 is marginally lower at 0.1 μM ATP than at 1 μM ATP. These tendencies cannot be explained by the energetics of M5. The helical pitch of F-actin remains virtually unchanged during the Brownian search by the detached head, as judged from the foot sliding rate (Table 2) that reflects the frequency of changes in the helical pitch of F-actin. Therefore, even when taking the MD-separation of *N*_m-m_ = 15 is energetically unfavourable, M5 always binds to the corresponding actin site. One possibility that explains these nucleotide-dependent tendencies is that the structure of phalloidin-bound F-actin may be susceptible to modulation by nucleotides in solution, in an unknown mechanism. This possibility is suggested by the nucleotide dependence of the foot sliding rate (Table 2).

**Table 2.** Foot sliding events observed under three nucleotide conditions.

| Nucleotide conditions | Total number of images | Number of foot sliding events | Imaging rate (fps) | Foot sliding rate ($s^{-1}$) |
| --- | --- | --- | --- | --- |
| 1 mM ADP | 25,790 | 287 | 3.0 | 0.033 |
| 0.1 μM ATP | 7,441 | 135 | 6.67 | 0.12 |
| 1 μM ATP | 5,709 | 175 | 6.67 | 0.20 |

## Discussion

Before discussing the energy conversion mechanism in M5 walking, it is important to recapitulate the recent debate on the timing of Pi release in myosins^28–34,36,59^. The debate is now being settled, to which the crystal structure of myosin VI has contributed greatly^29^. This structure shows the presence of Pi in the backdoor tunnel^35^, not in the active site, while the lever-arm is in the pre-powerstroke state^29,36,59^. Therefore, this structure suggests the possibility that the powerstroke occurs when Pi is still somewhere near the tunnel exit or at the tunnel exit, thereby explaining the observations of ∼2-times slower release of Pi into solution than the lever-arm swing^31–33^. A structural comparison between the primed pre-powerstroke state and the post-powerstroke state, as revealed by time-resolved cryo-EM, also supports this view^23^. It is hard to believe that a large force can be generated in weakly bound myosin head with ADP·Pi at the active site, without any chemical transition. We therefore adopt the most plausible view that the force generation occurs at some time after Pi has left from the active site but before it is released into solution. However, even in this view, we will simply describe the force generation as occurring after Pi release.

Despite numerous single-molecule studies of M5 motility using optical methods^9–12,15,17,18,20,25,47.48,50–53^, the ATP energy conversion in M5 walking has been little studied. The energy conversion mechanism in M5 is derived from the prevailing view developed from extensive studies of the muscle actomyosin system^1,2,27,36,60^. Changes in the ATP free energy along the muscle actomyosin ATPase reaction pathway have been estimated from measurements of ATPase kinetics and thermodynamics^61,62^. In this system, the free energy changes occur primarily at the stages of ATP binding-induced head dissociation^19^, ATP hydrolysis to ADP·Pi accompanied by a recovery stroke^20–22^, Pi release accompanied by the change from weak to strong head binding to actin^29,36^, and ADP release^17,37^. The amounts of released energy in these stages are approximately 10, 2, 12, and 3 *k*_B_*T*, respectively^61^ (see Fig. 5a). This profile of free energy changes is likely to be similar in the M5−actin system, although the free energy changes at the first (ATP binding), fourth (Pi release), and last (ADP release) stages depend on the concentrations of free ATP, Pi, and ADP, respectively. Indeed, the actin-activated hydrolysis of ATP to ADP·Pi by M5 is associated with a free energy change of −2.3 k_B_T^16^, similar to the case of muscle myosin. As contractile force is generated when muscle myosin is in the strongly bound state formed during or after Pi release, it is rational to assume that the 12 kBT of free energy provided at the stage of Pi release is used to rotate or strain the crossbridge at the converter hinge^1,2,27^; when this idea is applied to M5, this process may be delineated as winding up a spiral spring at the converter hinge of the L-head (Fig. 5b), as structurally suggested recently^23^. It has therefore generally been thought that the chemical energy of ATP is the source of the mechanical work (or strain energy) produced by the power stroke. However, this energy conversion mechanism cannot account for the ATP-free walking observed in this study.

**Fig. 5.**
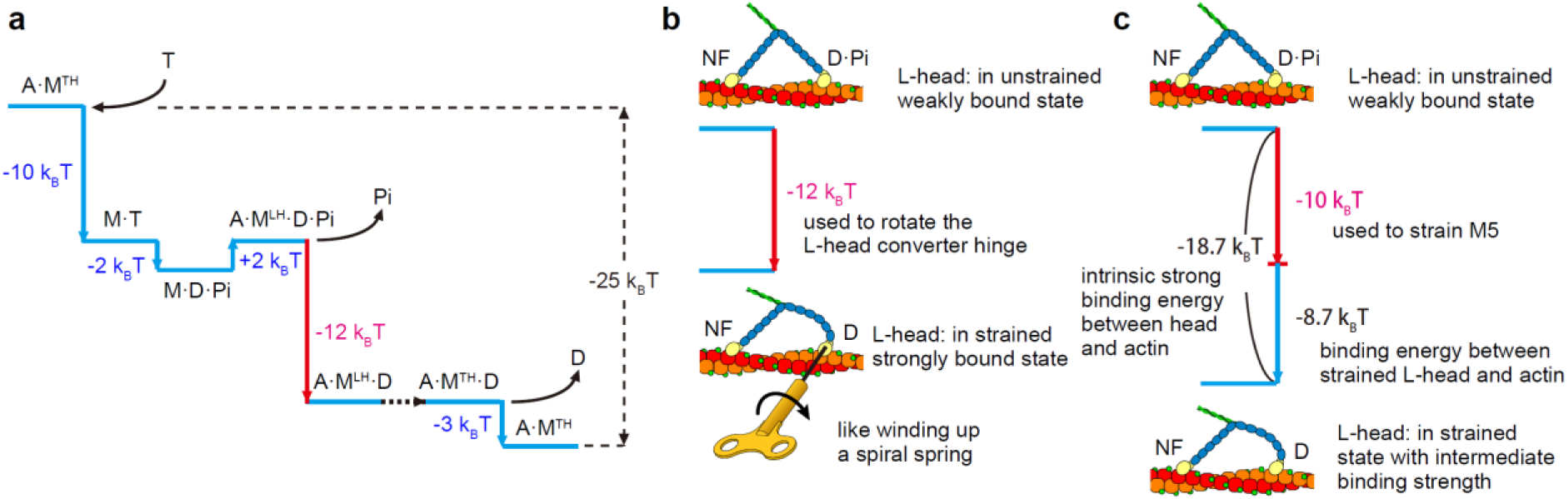
Mechanism of energy conversion into mechanical work in M5. A, M, T, D, Pi, and NF denote actin, M5, ATP, ADP, orthophosphate, and nucleotide-free, respectively. **a** Free energy changes along the ATPase reaction of M5−actin. Since some values of free energy changes are not yet available for M5−actin, the values shown correspond to those for skeletal actomyosin^61^. The superscripts, TH and LH, denote the trailing head and leading head, respectively. During the transition from A·M^LH^·D to A·M^TH^·D (black dotted line), the second head detaches from actin to become the new L-head. **b** Conventional mechanism of energy transduction to mechanical work. In this mechanism, the free energy ∼12 *k*_B_*T* released at the transition from A·M^LH^·D·Pi to A·M^LH^·D, shown with the red line in (**a**), is assumed to be used to actively rotate the lever-arm around the L-head converter hinge, like winding up a spiral spring. By this rotation, the L-head is strained. **c** New mechanism proposed in this study. More than 50% of the strong binding energy between actin and the ADP-bound L-head is used to strain the M5 molecule predominantly at the L-head. This results in weaker binding of the L-head to actin. Here, the binding energy between M·D·Pi and actin is assumed to be negligibly small, compared to that between M·D and actin.

In this study, we have presented two main lines of data supporting our view that the strong binding energy between actin and ADP-bound M5 is converted into mechanical work during ATP-dependent walking (Fig. 5c). Note that in our assay system, a load is applied against the walking of M5 from the substrate surface. The first line of data shows that the strain energy accumulated in the L-head, *U*_LCH_(*φ*_L_) + *U*_neck_ [*h*(*φ*_L_)], is almost identical between ATP-free and ATP-dependent walking for the respective cases of MD-separations (Fig. 3e & Supplementary Fig. 6e,f). It is difficult to understand this good quantitative agreement of strain energy if the mechanism of strain energy generation and hence the mechanism of energy conversion into mechanical work differed significantly between ATP-free and ATP-dependent walking. Therefore, this good agreement strongly suggests that during the ATP-dependent walking of M5, the chemical free energy of ATP is not converted into the strain energy of ∼10 *k*_B_*T* and is therefore not converted into mechanical work. This view may not be far from the traditional view that muscle contractile tension is generated when actin−myosin binding is strengthened by the release of Pi, which is accompanied by the release of free energy by 12 *k*_B_*T*. It is impossible (at least very difficult) to decompose this released energy into its individual components, such as the free energy changes caused by the Pi release itself, by binding between actin and ADP-bound head, and by conformational changes in myosin and actin. However, a significant fraction of the released free energy must originate from the strong head−actin binding energy.

The second line of supporting data is the foot stomping (brief head detachment) that occurs predominantly at the L-head in all the nucleotide conditions tested. This spontaneous brief detachment can be understood as being caused by the reduction in head-actin binding energy. The foot stomp rate was found to be comparable, irrespective of the different nucleotide conditions tested, thereby indicating a comparable level of reduction in head-actin binding energy, corresponding to the strain energy of ∼10 *k*_B_*T*. In addition to the two lines of data mentioned above, the energetic analysis and the computational simulations of ATP-free walking could quantitatively explain the success/failure ratio for the forward step of M5 caused by mechanical detachment of the T-head. Our proposition that the L-head−actin binding energy serves as the energy source for mechanical work appears to imply that the magnitude of the force generated by a single M5 molecule in the presence of ATP depends on the ionic strength; the force becomes greater as the ionic strength is reduced. However, the molecular strain of M5 in the DHB state does not seem sensitive to the ionic strength, as far as the L-head−actin binding is strong enough. In muscle, it is well known that the isometric tension increases as ionic strength decreases^e.g. 63,64^. However, interpreting this increased force in muscle is not straightforward. Single-molecule force measurements are ideal for addressing this issue. To the best of our knowledge, however, systematic measurements of the force generated by a single myosin molecule have not been performed at different ionic strengths.

Our view that M5 does not convert ATP chemical energy into mechanical work does not mean that this chemical energy is not essential for the forward translocation. To use the strain energy generated by L-head−actin binding for forward translocation, the strong association between the T-head and actin must be broken^19^. ATP binding to the T-head breaks this strong bond, releasing free energy of approximately 10 kBT, assuming that it is approximately the same between muscle myosin and M5. Although it is very difficult to decompose this released free energy into constituent components, the large binding energy between the NF T-head and ATP must be used to break the T-head−actin bond. The binding energy between the NF T-head and actin is approximately 23.8 *k*_B_*T*, corresponding to the equilibrium constant of 4.9 × 10^−11^ M (Ref. 16). Although the affinity of the NF head for ATP is too high to measure experimentally, the ATP binding energy must be comparable to 23.8 *k*_B_*T*. In addition, the hydrolysis of ATP to ADP·Pi associated with a recovery stroke^20–22^ and a free energy change of −2.3 *k*_B_*T* (Ref. 16) plays an important role in increasing the efficiency of the forward landing of the detached head as well as in preventing its backward return. Thus, the major role of ATP chemical energy is to release the strain energy acquired by the strong L-head−actin binding for the forward movement of M5. In addition, it lowers the energy barrier for the forward step and raises the energy barrier for the backward return, thereby ensuring the unidirectional process of M5 walking.

It is well known in various contexts that a protein can change its affinity for ligands, membranes, or partner proteins when subjected to external mechanical stress^e.g.,65–67^. However, it has only rarely been reported that a homo-oligomeric protein changes its binding affinity for a target as a result of the structural strain caused by the multivalent binding. For example, double-headed IgGs were observed to migrate non-unidirectionally on a two-dimensional antigen lattice, when the adjacent epitopes were appropriately separated, whereas single-headed IgGs stably remained at one antigenic site^68^. There are many homo-oligomeric proteins with structural symmetry that can bind to a target with multiple binding sites^69^. Although not yet explored, these oligomeric proteins may universally exploit the affinity reduction caused by strain generation due to multivalent binding. M5 may be a special case where the strain energy gained in this way is used for unidirectional movement and mechanical work. This energy conversion mechanism will be found in other motor proteins and used for the potential creation of novel molecular motors.

## Methods

### Sample preparation

The full length of M5 was prepared from chick brain^70^ and digested with proteinase K in the absence of Ca^2+^ to produce M5-HMM^39,41^. Instead of using an actin-copelleting method to purify M5-HMM^39,41^, we used a protein concentrator (Vivaspin 500 of 100,000 MWCO; Sartorius) and washing buffer containing 20 mM imidazole-HCl (pH 7.6), 100 mM KCl, 2 mM MgCl_2_, 1 mM EGTA, 5 mM DTT and 0.1 mM PMSF. This was because the short coiled-coil tail of M5-HMM occasionally unfolds when it is bound to actin filaments in the absence of ATP^38,39^. Partially biotinylated (20%) actin filaments from rabbit skeletal muscles were prepared and then stabilized with phalloidin^10^. Streptavidin, glucose, hexokinase and nucleotides were purchased from Wako Pure Chemicals, and phalloidin was purchased from Sigma Aldrich. The synthetic lipids, 1,2-dipalmitoyl-sn-glycero-3-phosphocholine (DPPC), positively charged 1,2-dipalmitoyl-3-trimethylammonium-propane (DPTAP), and 1,2-dipalmitoyl-sn-glycero-3-phosphoethanolamine-N-(cap biotinyl) (biotin-cap-DPPE), were purchased from Avanti Polar Lipids.

### HS-AFM observations

A mica disc (1 mm in diameter) was glued onto the top of a sample stage (glass rod of 2 mm in diameter and 2 mm in height) and its top surface was cleaved using Scotch tape. The mica-supported lipid bilayers as a substrate surface were formed by the deposition of liposomes prepared using DPPC, DPTAP and biotin-cap-DPPE (with a typical weight ratio of 0.85:0.05:0.1) onto the freshly cleaved mica surface^38,71^. After rinsing the lipid bilayer substrate with buffer A (20 mM imidazole-HCl (pH 7.6), 25 mM KCl, 2 mM MgCl_2_, 1 mM EGTA, 5 mM DTT), a drop (2 μl) of streptavidin in buffer A (10 nM) was deposited on the surface for 3 min. After rinsing with buffer A, a drop (2 μl) of partially biotinylated actin filaments (1 μM) in buffer A was deposited on the substrate surface for 10 min. After rinsing with a solution containing 1 mM ADP (1 U ml^−1^ hexokinase and 10 mM glucose were added to remove contaminating ATP) in buffer A, we deposited a drop (2 μl) of the same solution plus M5-HMM (0.1 to 1 nM) on the substrate surface and allowed it to sit for 3 min. Finally, the sample stage was mounted onto the scanning stage of our HS-AFM apparatus^72,73^ (to which the interactive mode was newly implemented) and immersed in a cell chamber filled with the same solution (60 μl) without M5-HMM. All HS-AFM imaging experiments were performed at room temperature (25°C) in amplitude modulation mode. For the usual imaging (no sample manipulation by the AFM tip) of the M5-HMM interacting with actin, the cantilever free oscillation amplitude *A*_0_ was set to 1.5 nm, while the set point amplitude during imaging *A*_s_ was set to 0.85 × *A*_0_ (= 1.28 nm). The mean oscillation energy loss per tip‒sample contact, *ΔE*_ts_, was estimated to be *ΔE*_ts_ = γ*k*_c_(*A*_0_^2^ ‒ *A*_s_^2^)/(2*Q*_c_) = 3.1‒4.7 *k*_B_*T* (*k*_B_, Boltzmann constant; *T* = 298 in kelvin), where γ (≈ 0.9) is a correction factor that accounts for the amplitude reduction caused by a cantilever resonant frequency shift toward a higher frequency by repulsive tip‒sample interactions, and *k*_c_ = 0.1‒0.15 N/m and *Q*_c_ = 1.5 are the spring constant and the quality factor of the short cantilevers used. For interactive mode imaging, the set point amplitude *A*_p_ was typically set to 0.33 × *A*_0_ (= 0.5 nm) to apply a strong force to a few closely adjacent positions on either head of the actin-bound M5-HMM. Short cantilevers (BL-AC7DS-KU4; 7−8 μm long, 2 μm wide and 90 nm thick) were custom-made by Olympus, whose resonant frequency, spring constant and quality factor were *f*_c_ = 0.8−1.2 MHz in water. The spring constant of each cantilever was estimated from its measured frequency spectrum of thermal oscillation in buffer A. The interactive mode is described in Supplementary Fig. 1.

### Caged-ATP experiments

For caged-ATP experiments combined with iHS-AFM observations, we installed the following optical components in our HS-AFM system (Supplementary Fig. 5a,b): a UV-LED system of 365 nm (L11922-401 and C11924-111, Hamamatsu photonics, Japan) and a dichroic mirror that reflects UV light (320‒360 nm) and transmits visible light (500‒700 nm). Continuous UV light was irradiated onto a sample area around the cantilever through the same objective lens (ELDW 20XC, Nikon, Japan) used for cantilever deflection detection. The power of the incident UV light at the exit of the objective lens was 19 mW at 350 nm. The diameter of the focal region measured by the half-intensity diameter was approximately 300 μm (Supplementary Fig. 5c), from which the focal volume was estimated to be ∼14 nL. As the volume of the liquid cell was 60 μL, the caged-ATP experiments could be repeated many times. After M5-HMM was deposited onto a mica-supported lipid bilayer surface containing immobilized actin filaments, the sample stage was rinsed with a solution containing 20 μM caged-ATP, 1 U ml^−1^ hexokinase, and 10 mM glucose dissolved in buffer A. Therefore, the ATP released from caged-ATP was quickly converted to ADP. Finally, the sample stage was immersed in a cell chamber filled with the same solution (60 μl) without M5-HMM. Before performing iHS-AFM observations, photolysis of caged-ATP was performed several times to confirm whether M5 moved unidirectionally by photolysis of caged-ATP. This experiment generated some ADP in solution, and therefore, the M5 molecules in the observation area were presumably bound to ADP before the subsequent iHS-AFM imaging. Then, ATP-free walking was observed for several steps, immediately after which caged-ATP was photolyzed while HS-AFM images were captured continuously (Supplementary Fig. 5d; Supplementary Video 3).

### Mechanical properties of converter hinge in actin-bound single-headed M5

The potential energy *U*(*φ*) of the converter hinge determines the equilibrium distribution *P*(*φ*) ∝ exp (−*U*(*φ*)/*k*_*B*_*T*) of the lever-arm angle *φ* relative to the orientation of F-actin measured from the plus end side (Fig. 3a, inset). By fitting to the HS-AFM data of single-headed M5 bound to actin observed in 1 mM ADP and NF conditions^38^, we found that in both conditions, *U*(*φ*) fits well to the functional form of *U*(*φ*) = [(*bk*_0_ + *k*_1_(*φ* − *φ*_0_))/(*b* + (*φ* − *φ*_0_))](1 − cos(*φ* − *φ*_0_)) (Fig. 3b, black lines). Here, *φ*_0_ is the equilibrium angle: 28.5° at 1 mM ADP and 32.9° in the NF condition. The other parameters are *b* = 34.4°, *k*_0_ = 23 *k*_B_*T* and *k*_1_ = 2.3 *k*_B_*T* at 1 mM ADP, and *b* = 75.0°, *k*_0_ = 37.9 *k*_B_*T* and *k*_1_ = 4.1 *k*_B_*T* in the NF condition. The stiffness of the converter hinge in 1 mM ADP and NF conditions, i.e., the angular spring constants, *K*_D_(*φ*) and *K*_NF_(*φ*), were determined from *U*(*φ*) = (1⁄2) *K*(*φ*) · (*φ* − *φ*_0_)^2^ (Fig. 3b, red lines). The magnitude of the force *F*(*φ*) in the direction parallel -to F-actin generated at the free end of the lever-arm was obtained as *F*(*φ*) = *K*(*φ*) · (*φ* − *φ*_0_)^2^/[*L*_neck_(*cosφ*_0_ − *cosφ*)]/0.243 (Fig. 3b, blue lines), where *L*_neck_ is the neck length (∼23 nm estimated from nsEM images^42^) and the division by 0.243 is for unit conversion from *k*_B_*T* to pN·nm.

### Strain energy of M5 in DHB state

To estimate the strain energy accumulated in a M5 molecule in the DHB state, we first measured the following three geometrical parameters from HS-AFM images (Fig. 3c, inset): the tangential angle of the L-head’s neck region emerging from the motor domain, *φ*_L_; the angle of the T-head neck as a function of *φ*_L_, *φ*_T_(*φ*_L_); the deflection of the neck end of the L-head as a function of *φ*_L_, *h*(*φ*_L_). The angles, *φ*_L_ and *φ*_T_, relative to the orientation of F-actin, were measured from the plus end side of F-actin. The results are shown in Fig. 3c,d (for *N*_m-m_ = 13 actin subunits), Supplementary Fig. 6a,c (for *N*_m-m_ = 11 actin subunits), and Supplementary Fig. 6b,d (for *N*_m-m_ = 15 actin subunits). The total strain energy *E*_total_ is expressed as *E*_total_ = *U*_D_(*φ*_L_) + *U*_X_[*φ*_T_*(φ*_L_)] + *U*_neck_[*h*(*φ_L_*)], where *U*_D_(*φ*_L_) is the strain energy of the L-head converter hinge (the subscription D denotes the ADP-bound state), *U*_X_[*φ*_T_*(φ*_L_)] is the strain energy of the T-head converter hinge (the subscription X is either D or NF depending on the nucleotide state of the T-head), and *U*_neck_[*h*(*φ_L_*)] is the bending energy of the L-head’s neck region. The DHB states formed during ATP-free and ATP-dependent walking are almost stationary, meaning that all forces acting on a M5 molecule, including constraining forces, are balanced. From the principle of ‘virtual work’ in mechanics, the work done by the non-constraining forces (i.e., the forces generated by the two converter hinges and the bending of the L-head) is zero, which can be expressed as

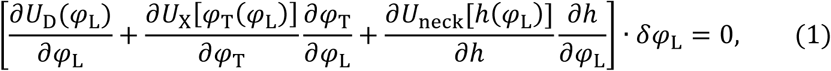

where *δφ*_L_ is the virtual displacement. Therefore, the total strain energy *E*_total_ is conserved. The results obtained for *E*_total_, *U*_D_(*φ*_L_) + *U*_X_[*φ*_T_*(φ*_L_)], *U*_neck_[*h*(*φ_L_*)], *U*_X_[*φ*_T_*(φ*_L_)], and the spring constant of the L-head neck *k*_neck_ are shown in Fig.3e,f for *N*_m-m_ = 13, Supplementary Fig.6e,g for *N*_m-m_ = 11, and Supplementary Fig.6f,h for *N*_m-m_ = 15.

### Mechanical model and simulations

To obtain relevant numerical estimates for ATP-free stepping from mathematical modelling, a simple representation of the M5 dimer was employed, with the shape (in the characteristic HS-AFM planar geometry) characterized by only three variables: the angles *ϕ*_1_and *ϕ*_2_ for the two neck–motor domain junctions and the neck–neck junction angle *ϕ*_*c*_ (Supplementary Fig. 9). To simplify the simulations, our energy does not include a contribution due to elastic bending of the necks (i.e., *h* = 0). Instead, this elastic energy was included to the energy of L-head converter hinge. The neck–motor domain junction angle *ϕ* used in the M5 model and the maximum lever-arm angle 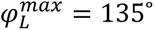 are related via 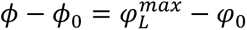 where the equilibrium angle *ϕ*_0_ = 148.5° of the neck–motor domain junction corresponds to the equilibrium lever-arm angle *φ*_0_ = 28.5° relative to the actin filament (see Supplementary Methods).

The total energy of DHB M5 is *E* = *U*(*ϕ*_1_) + *U*(*ϕ*_2_) − 2*∈*, where *∈* is the absolute value of the actin-binding energy of the M5 head. As done for previous models^54,55^, we assumed that the neck– neck junction was highly flexible and that there was no contribution to the elastic energy coming from it.

Conformational motions that underlie M5 stepping proceed on the timescales of milliseconds, where inertial effects are negligible, and only overdamped motions with velocities instantaneously following the applied forces occur. Therefore, the stochastic Langevin description, which effectively accounts for the interactions of M5 with the surrounding environment via viscous friction, is applicable. To numerically investigate ATP-free M5 translocation, Brownian dynamics simulations were performed for the five beads representing the rigid rods in our mechanical model (Supplementary Figs. 9,10). The dynamic evolution of the M5 five-bead model is therefore described by equations for the beads 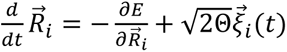, 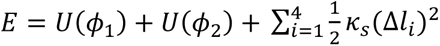 and 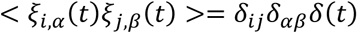 Here, 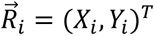 is the instantaneous position of bead *i*. In the reduced equations, lengths were measured in units of the neck length 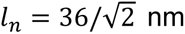, and time was measured in units of 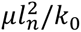, where *μ* is the viscous friction coefficient of beads and *k*_0_ = 23 *k*_*B*_*T*. Hence, converter hinge energies *U*(*ϕ*) were measured in units of *k*_0_. The total potential energy *E* of the 5-bead model is the sum of both converter hinge energies. The expression of *E* contains additional barriers of dimensionless strength ***κ***_*s*_ = 150, which prevent length changes Δ*l* of the motor domains and neck rods 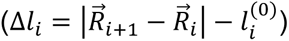. The dimensionless quantity Θ, which determines the fluctuation intensity of beads, was related to the equilibrium converter hinge stiffness *k*_0_, i.e., Θ = *k*_*B*_*T*/*k*_0_ = 1/23. Starting from the configuration with both myosin heads bound to the actin filament, the set of equations was numerically integrated to obtain bead positions 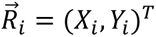at any time moment. In the simulations, the Euler scheme with a dimensionless time-step of 5 · 10^−4^ was employed. In our dimensionless numerical model, time was measured in units of 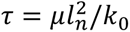. Introducing the bead mobility Γ = 1/*μ* and using the dimensionless quantity Θ = *k_B_T*/*k*_0_, we obtain 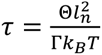. Using the Einstein relation Γ*k*_*B*_*T* = *D*, it is expressed as 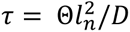. Assuming a diffusion constant of *D* = 10^−7^cm^2^/s, we obtain a time unit of approximately *τ* = 3 *μs*. This estimate was applied to relate timescales of model simulations to observations from HS-AFM experiments. Consistent with the approximate character of the model, the actin filament and interactions of M5 heads with it were accounted for in a simplified way. For details of the model, its parameters, and numerical simulations, refer to the Supplementary Fig. 9, Supplementary Table 1, and Supplementary Methods.

## Data availability

Data supporting the findings of this study are included within the article and supplementary figures. The videos analysed to generate Fig. 3c-f are partially included as Supplementary Videos 1 and 2 (because many video images were used for Fig.3c-f, we will provide them upon request). It is also the same for the videos analysed for Tables 1 and 2. Source data for Fig.3a,c,d, Fig. 4d-g, Supplementary Fig,6,a-d, and Supplementary Fig.10l are provided with this paper.

## Code availability

This study does not produce code applicable for other purposes.

## Acknowledgements

We thank A. Mikhailov for advising us about the modelling and simulations of ATP-free walking of M5, and R. Vale and J. Howard for critical reading of the original manuscript. We acknowledge Grant-in-Aid for Scientific Research (S) #24227005 and #17H06121 (T.A.), (A) #22H00405 (T.A.) and (C) #21K03483 (H.F.) from the Japan Society for the Promotion of Science, the Human Frontier Science Program grant #RGP0019 (T.A.), the PRESTO grant #JPMJPR13L4 (N.K.) from the Japan Science and Technology Agency, and the Transdisciplinary Research Promotion grant (H.F. and N.K.) from NanoLSI-WPI for financial support of this work.

## Author information

### Contributions

T.A. conceptualized this study and the iHS-AFM technique. N.K. constructed a UV-illumination system and installed it to the HS-AFM system, performed all experiments and analysed the image data. T.U. developed a software program for iHS-AFM. H.F. performed modelling and simulations of ATP-free M5 translocation. T.A. analysed the strain energy. H.F. and T.A. analysed the energetics of ATP-free and ATP-dependent M5 walking. T.A. wrote the manuscript and all authors provided feedback and editorial support.

Correspondence and requests for materials should be addressed to Toshio Ando

## Additional Information

Supplementary Information

Description of Additional Supplementary Files

