## Supplementary Figures 1-10 for "Energy conversion mechanism revealed by ATP-free and ATP-dependent walking of myosin V motor"


| | |
|---|---|
| Supplementary Fig. 1 | Interactive HS-AFM (iHS-AFM) |
| Supplementary Fig. 2 | Directional rule of M5 walking along actin filaments |
| Supplementary Fig. 3 | iHS-AFM images showing various head detachment events |
| Supplementary Fig. 4 | iHS-AFM images showing the stepping direction independent of the scanning direction |
| Supplementary Fig. 5 | Caged ATP experiments showing ATP-dependent waking activity of M5 after repeated applications of strong force |
| Supplementary Fig. 6 | Mechanical properties of converter hinge and L-head neck of M5 with MD-separations of $N_{m\text{-}m}$ = 11 and 15. |
| Supplementary Fig. 7 | Foot stomping and foot sliding observed in the presence of 1 mM ADP |
| Supplementary Fig. 8 | Probability of taking different motor domain separations just after landing of detached T-head observed in different nucleotide conditions |
| Supplementary Fig. 9 | M5 dimer mechanical model |
| Supplementary Fig. 10 | Mechanical model and simulations of ATP-free walking along actin filaments |
| Supplementary Methods | Modelling of myosin V dimer; Modelling of actin filament; Modelling of interactions between M5 and actin filaments; Additional model constraints; Simulation course of ATP-free M5 stepping by T-head unbinding; |
| Supplementary Table 1 | Parameters and their values used for mechanical model and simulations of ATP-free M5 translocation; |
| Supplementary references | References cited in Supplementary Methods |

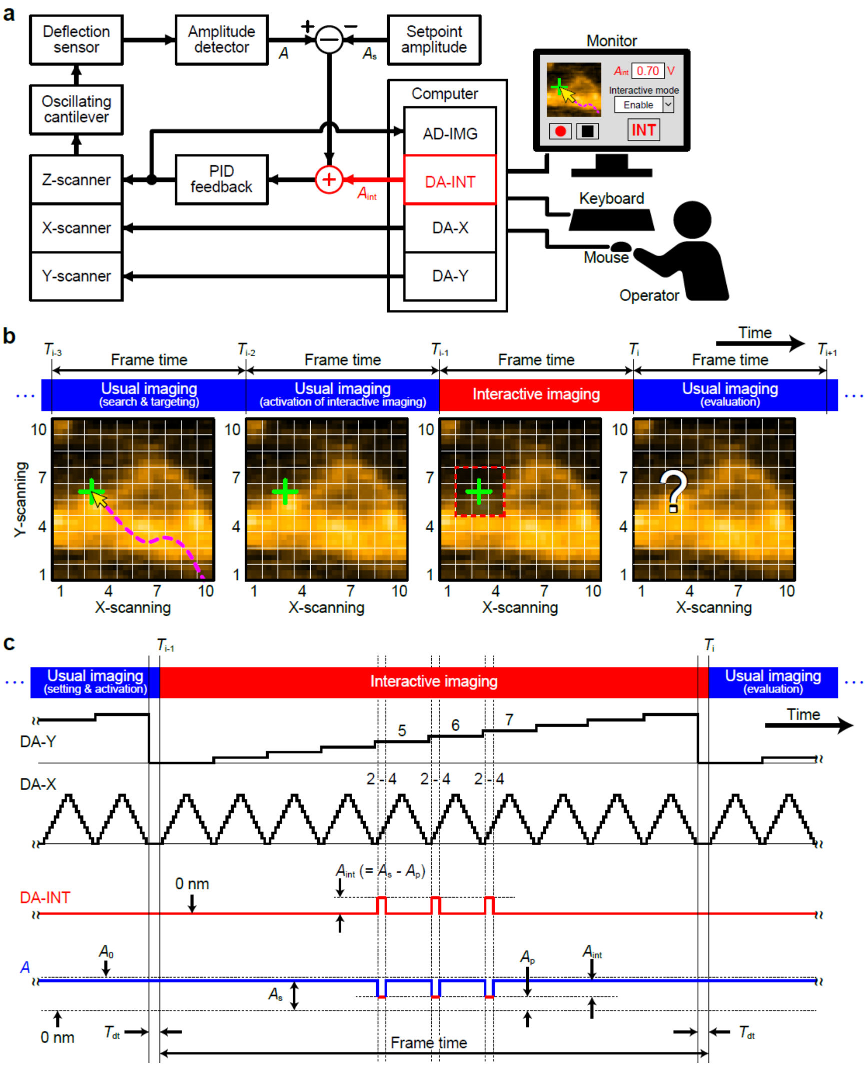


**Supplementary Fig. 1 | Interactive HS-AFM (iHS-AFM). a** Instrumental configuration for iHS-AFM. The iHS-AFM system was built by the addition of an adder circuit (red) and a DA board (DA-INT; red) to our HS-AFM system and by the modification of the operation software program. **b** Experimental procedure of iHS-AFM. Here, it is assumed that a strong force is applied to a region corresponding to the pixels of (2 + $m$, 5 + $n$) ($m$, $n$ = 0, 1, 2) in the image of 10 × 10 pixels. Before the interactive mode, the operator carries out the following procedures without halting the imaging: (i) setting the cantilever amplitude set point $A_p$, by which an excessive amplitude reduction, $A_{int} = A_s - A_p$, takes place during the interactive mode, (ii) searching a target area to which a strong force is to be applied, (iii) bringing a mouse pointer to the centre of the target area, and (iv) pushing a start key for the interactive mode. **c** Time diagrams before, during, and after the interactive mode. DA-X and DA-Y indicate output signals by which the scanner is displaced in the X- and Y-directions, respectively. DA-INT (red) indicates output signals by which the amplitude set point is reduced from $A_s$ to $A_p$ at the pixels of (2 + $m$, 5 + $n$) ($m$, $n$ = 0, 1, 2). $A$ (blue) indicates the amplitude signal. $T_{dt}$ indicates the time spent for data transfer from the AD board (AD-IMG) to the computer memory.

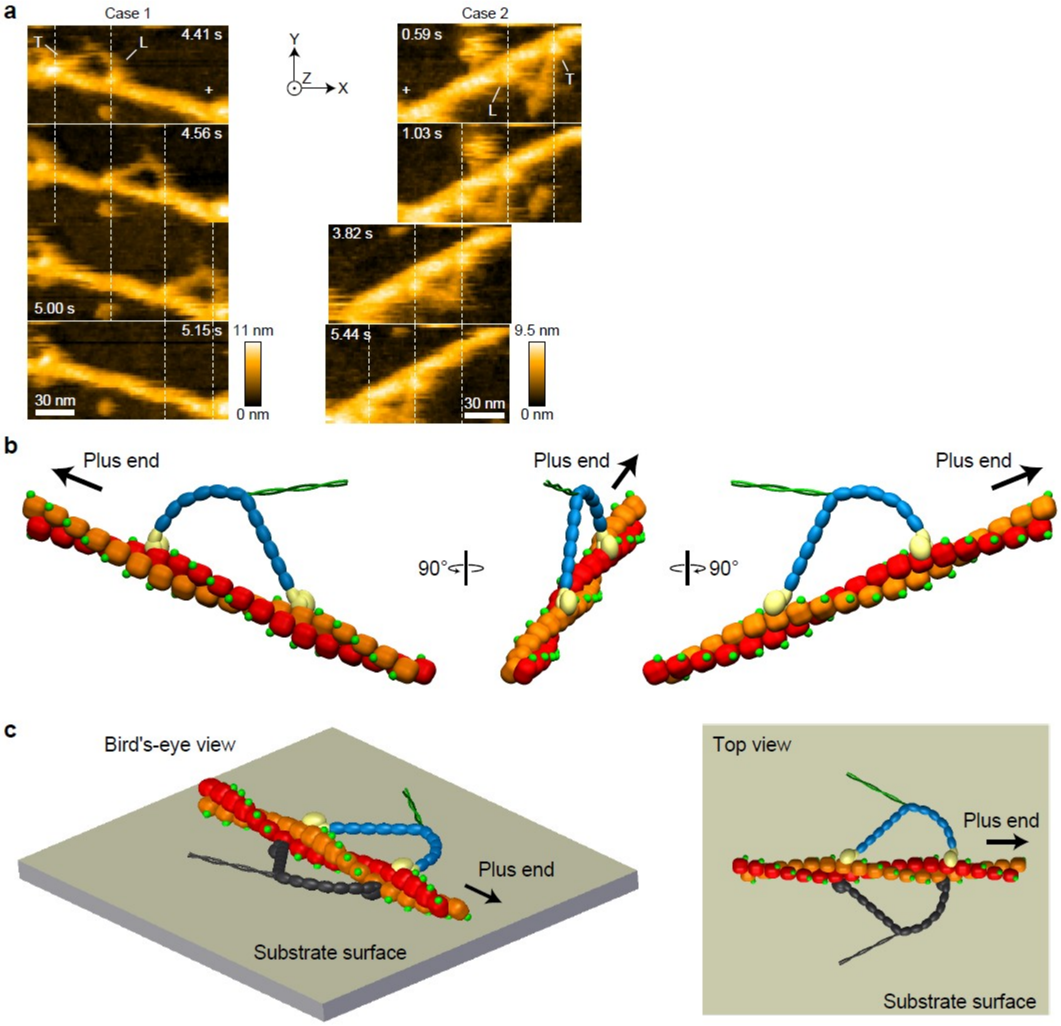


**Supplementary Fig. 2 | Directional rule of M5 walking along actin filaments. a** HS-AFM images showing unidirectional movement of M5 in the presence of 1 μM ATP. M5 moves from left to right (case 1) and from right to left (case 2). Vertical dashed lines, the centres of mass of the motor domains; plus sign, the plus end of actin filament; scan area, 155 x 75 $nm^2$ (case 1) and 141 x 76 $nm^2$ (case 2); number of pixels, 80 × 40; imaging rate, 6.7 fps. The images were acquired while the tip was being scanned towards the +X and +Y directions during raster scanning. Note that the motor domains look brighter than actin, meaning that the Z-height of the bound motor domains is larger than the top surface of the actin filament. **b** Cartoons showing M5 in the two-headed bound state from three angles. Note that the motor domains are bound to actin at their left flanks, consistent with the AFM images. **c** Cartoons showing a mechanism of the directional rule. (left) Bird's-eye view; (right) top view. The M5 in black does not follow the directional rule. Since its motor domains cannot enter the narrow space between the actin filament and the substrate surface, this manner of binding hardly occurs.

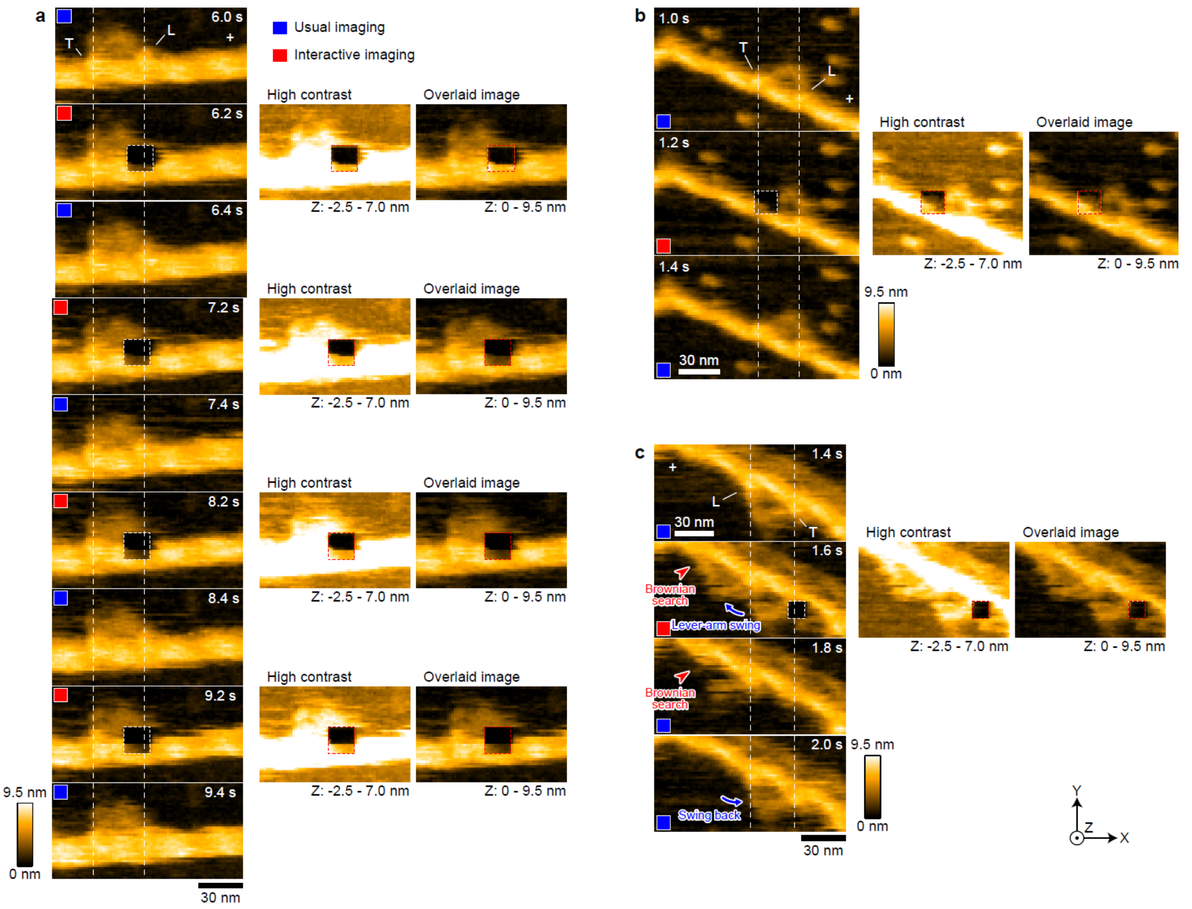


**Supplementary Fig. 3 | iHS-AFM images showing various head detachment events. a** Images showing events where the L-head detached by strong force application. Scan area, 130 × 65 nm$^2$; number of pixels, 80 × 40 pixels; imaging rate, 5 fps. A strong force was applied to the areas (11 × 11 pixels) marked by the white dashed lines (6.2, 7.2, 8.2 and 9.2 s). The high-contrast images were obtained by adjusting the brightness so that the brightness of the strong force applied regions became approximately identical to that of neighbouring regions in the corresponding left-hand images. The overlaid images were obtained by overlaying the high-contrast images of the strong force applied regions on the original images. The motor domain does not appear in the overlaid images, indicating that the motor domain was briefly detached and then returned to the original position as indicated in the images captured after the interactive mode. **b** Images showing events where the T-head did not detach by strong force application. Scan area, 150 × 90 nm$^2$; number of pixels, 80 × 48; imaging rate, 5 fps. A strong force was applied to the area (9 × 9 pixels) marked by the white dashed line (1.2 s). A motor domain appears in the overlaid image. **c** Images showing events where the T-head detached, after which its lever-arm swung forwards but returned to the original position (failure case). Scan area, 130 × 65 nm$^2$; number of pixels, 80 × 40; imaging rate, 5 fps. A strong force was applied to the area (7 × 7 pixels) marked by the white dashed line (1.6 s). The images were acquired while the tip was being scanned towards the +X and +Y directions during raster scanning.

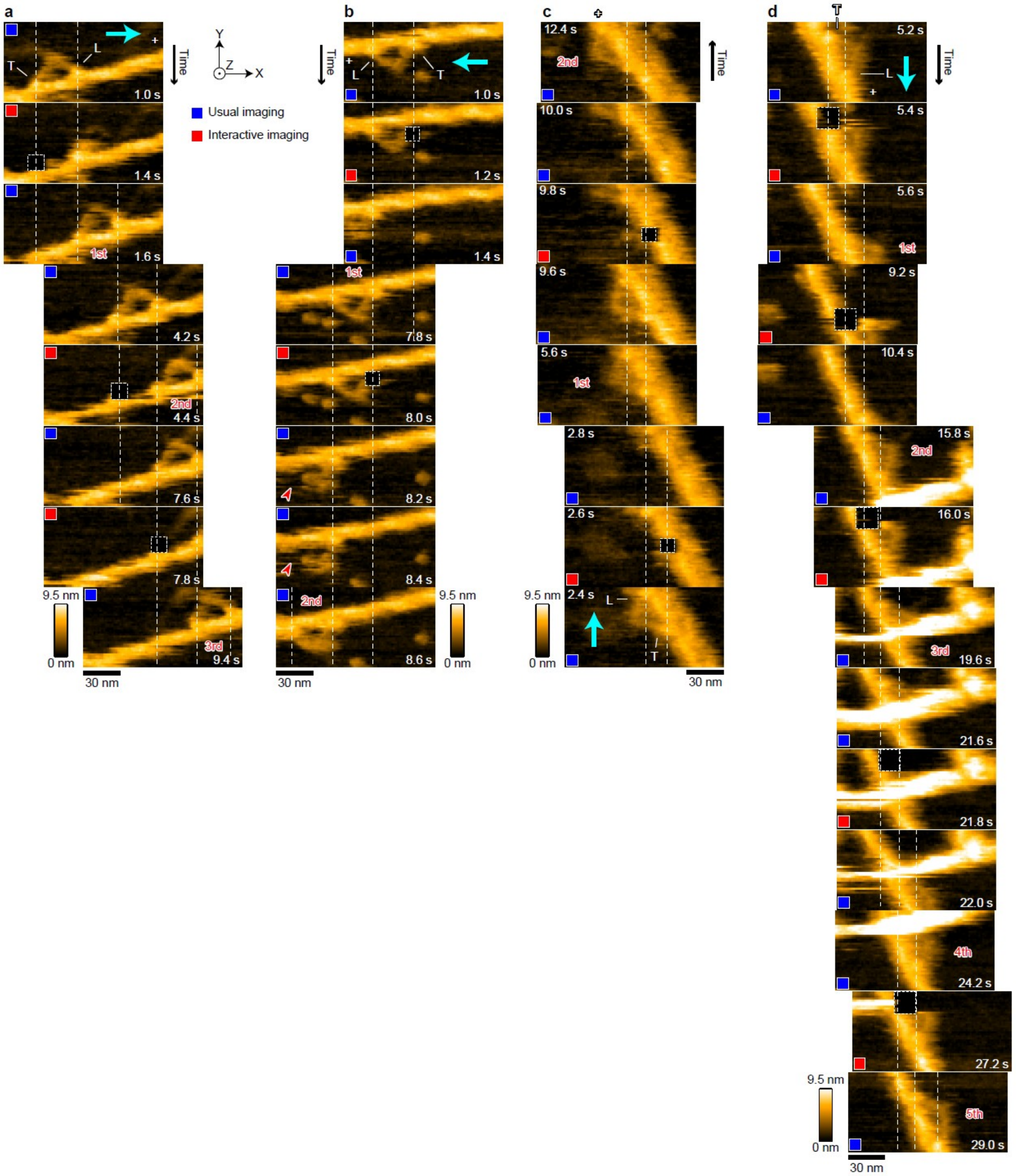


**Supplementary Fig. 4 | iHS-AFM images showing the stepping direction independent of the scanning direction. a** Stepping from left to right (3 steps), the same direction as that used for X-scanning during image capture. **b** Stepping from right to left (2 steps), opposite to the X-scanning direction during image capture. **c** Stepping from bottom to top (2 steps), the same direction as that used for the Y-scanning direction during image capture. **d** Long run (5 steps), stepping from top to bottom, opposite to the Y-scanning direction during image capture. Arrows in cyan, stepping directions; Scan area, 130 × 65 $nm^2$, number of pixels, 80 × 40; imaging rate, 5 fps; strong force applied area, 8 × 8 pixels for (**a**), 7 × 7 pixels for (**b** and **c**) and 11 × 11 pixels for (**d**). See Supplementary Video 2.

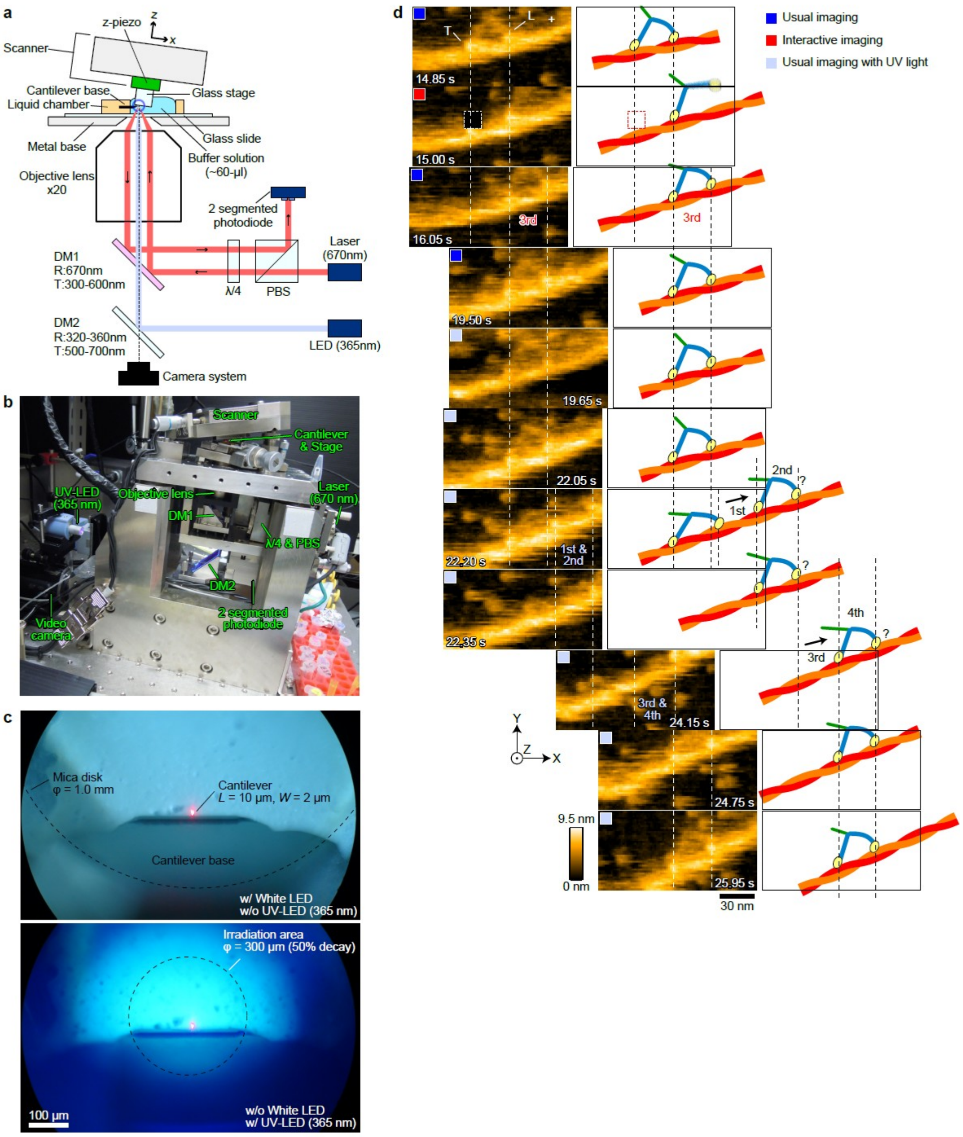


**Supplementary Fig. 5 | Caged ATP experiments showing ATP-dependent waking activity of M5 after repeated applications of strong force. a** Schematic showing the optical setup for the photolysis of caged-ATP installed in the HS-AFM head. A UV-LED (365 nm) was used for photolysis. DM, dichroic mirror; λ/4, a quarter-wave plate; PBS, polarizing beam splitter. **b** Picture showing the HS-AFM head with optics for the photolysis. **c** Optical microscope views around a cantilever without (top) and with (bottom) UV illumination. **d** Typical imaging result showing processive walking of M5 during photolysis of caged-ATP after iHS-AFM observations. The iHS-AFM observations were repeated three times before starting the caged-ATP photolysis from 19.65 s. Caged-ATP concentration, 20 μM. Scan area, 130 × 65 nm$^2$; number of pixels, 80 × 40; imaging rate, 6.7 fps. The solution contains hexokinase and glucose to quickly convert uncaged ATP into ADP so that the iHS-AFM and caged-ATP experiments can be repeated. Note that before the experiment shown in (**d**), the caged ATP experiments had been repeated so that the solution contained ADP to some extent. See Supplementary Video 3.

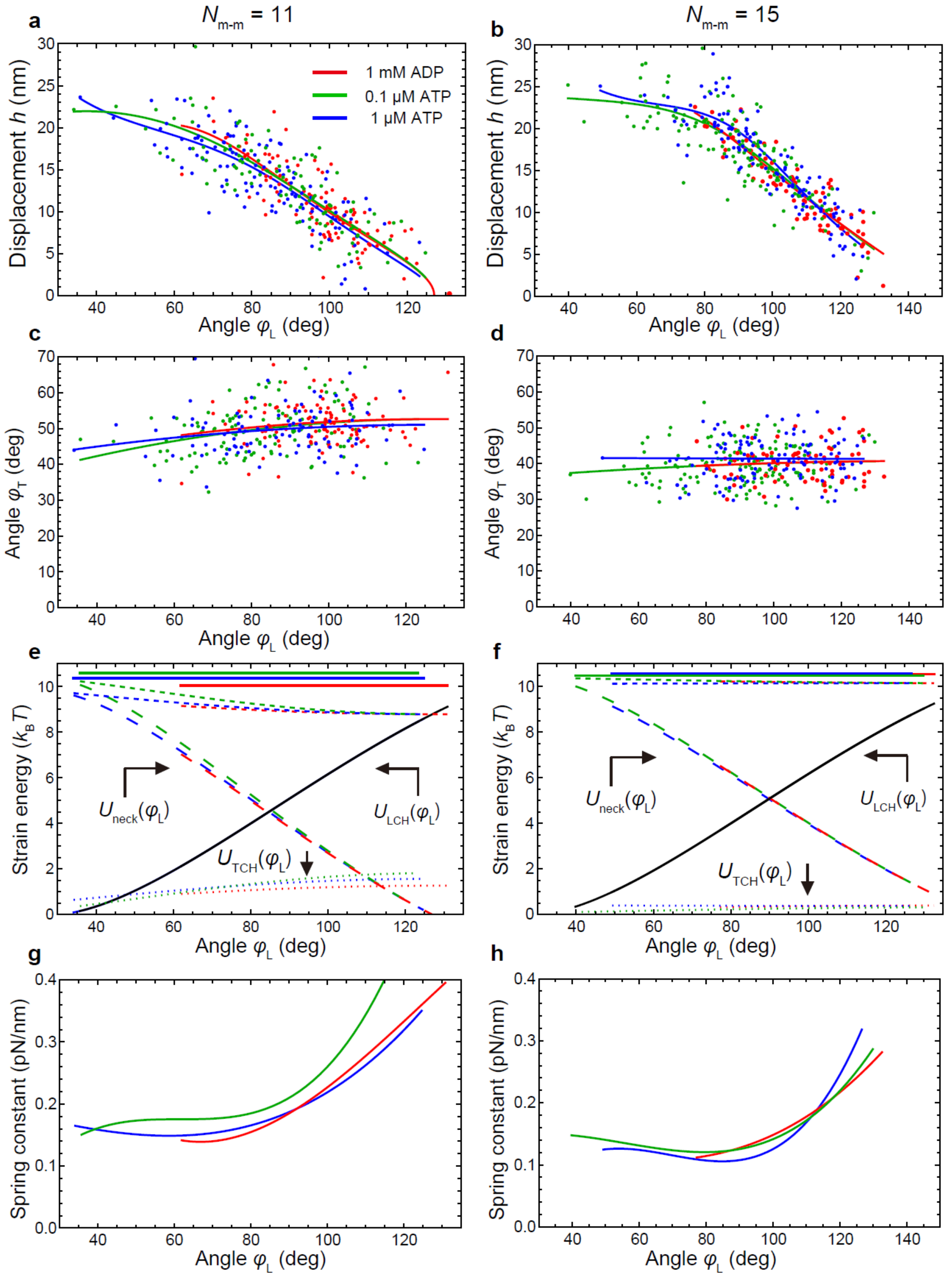


**Supplementary Fig. 6 | Mechanical properties of converter hinge and L-head neck of M5 with MD-separations of $N_{m\text{-}m}$ = 11 and 15. a,b** Lever-arm displacement depending on the angle $\varphi_L$. **c,d** Dependence of $\varphi_T$ on $\varphi_L$. **e,f** Strain energies at two converter hinges ($U_{TCH}$ and $U_{LCH}$) and L-head neck ($U_{neck}$). The top solid horizontal lines show total strain energy ($U_{TCH}$ + $U_{LCH}$ + $U_{neck}$), and the dashed lines slightly below the solid horizontal lines show $U_{LCH}$ + $U_{neck}$. **g,h** Spring constants of L-head neck bending, $k_{neck}$. The lines in (**a−f**) are drown within the observed angle rages of $\varphi_L$, except for the black lines in (**e,f**). The colour code shown in (**a**) is common in (**a−h**), except for the black lines in (**e,f**) identical in the three nucleotide conditions.

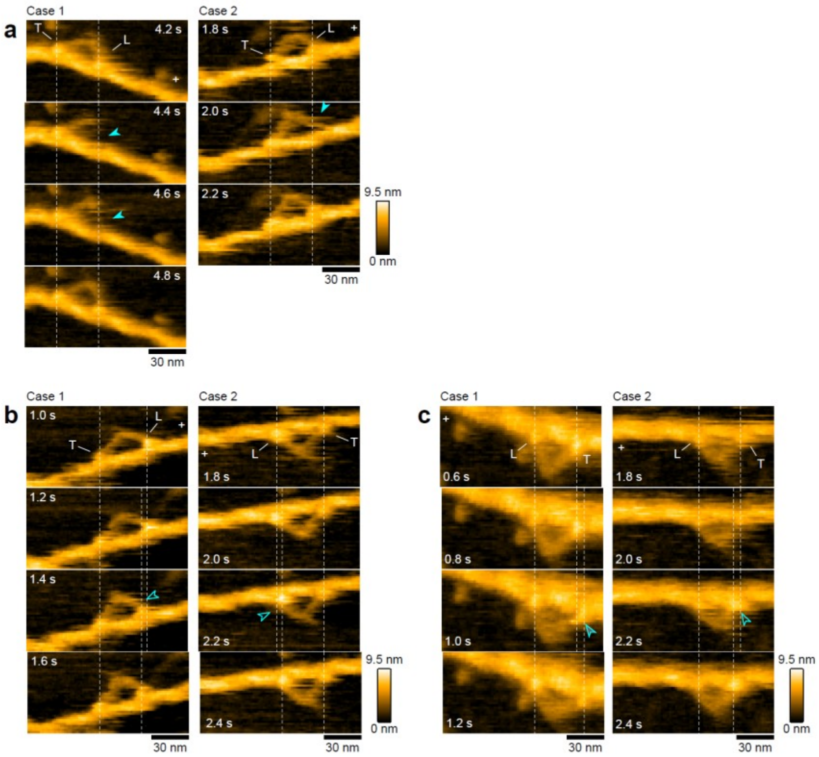


**Supplementary Fig. 7 | Foot stomping and foot sliding observed in the presence of 1 mM ADP. a** HS-AFM images showing foot-stomping events (cases 1 and 2) observed at the L-head. The arrows indicate the moments of L-head detachment (i.e., when the motor domain disappears from the images). See also Supplementary Video 4. **b,c** HS-AFM images showing foot sliding events at the L-head (**b**) and at the T-head (**c**). The motor domains pointed by the arrows are changing their position. See also Supplementary Video 5. The dashed vertical lines in **a−c** indicate the centres of mass of the motor domains.

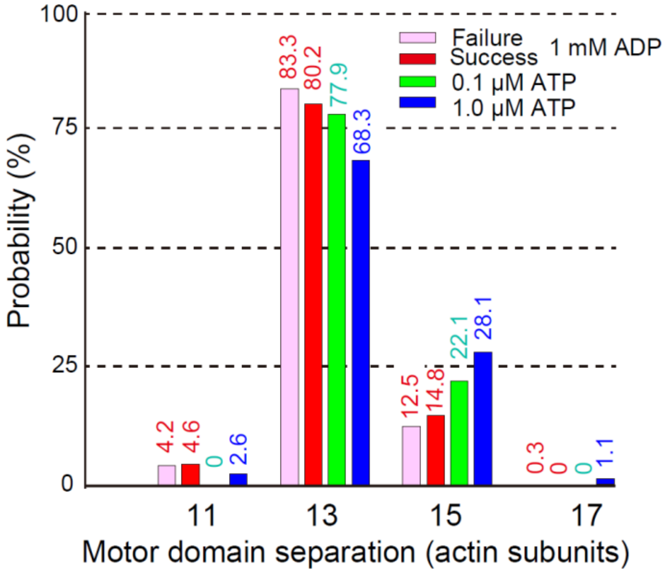


**Supplementary Fig. 8 | Probability of taking different motor domain separations just after landing of detached T-head observed in different nucleotide conditions.** 'Success' and 'failure' means that M5 steps forward or returns backward to an original actin or its vicinity after mechanical T-head detachment in the presence of 1 M ADP. Number of events: $n$ = 24 (1 mM ADP, failure); $n$ = 324 (1 mM ADP, success); $n$ = 68 (0.1 μM ATP); $n$ = 467 (1 μM ATP).

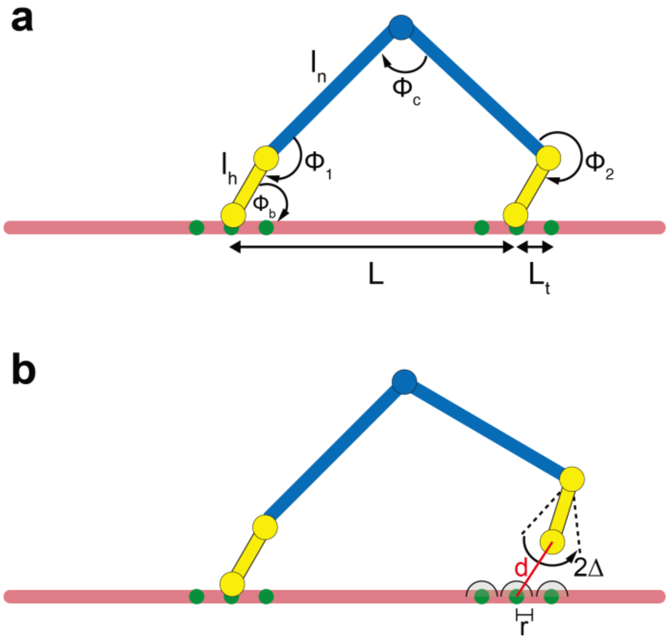


**Supplementary Fig. 9 | M5 dimer mechanical model. a** Rigid-rod representation of the M5 dimer and actin with notations of model variables and parameters. **b** Schematic illustration of conditions in which an unbound M5 head rebinds to actin. The tolerance window 2Δ for the relative orientation of the motor domain with respect to the fixed actin-bound orientation, the distance *d* from the motor domain to one of the actin binding sites, and the rebinding interaction radius *r* are indicated.

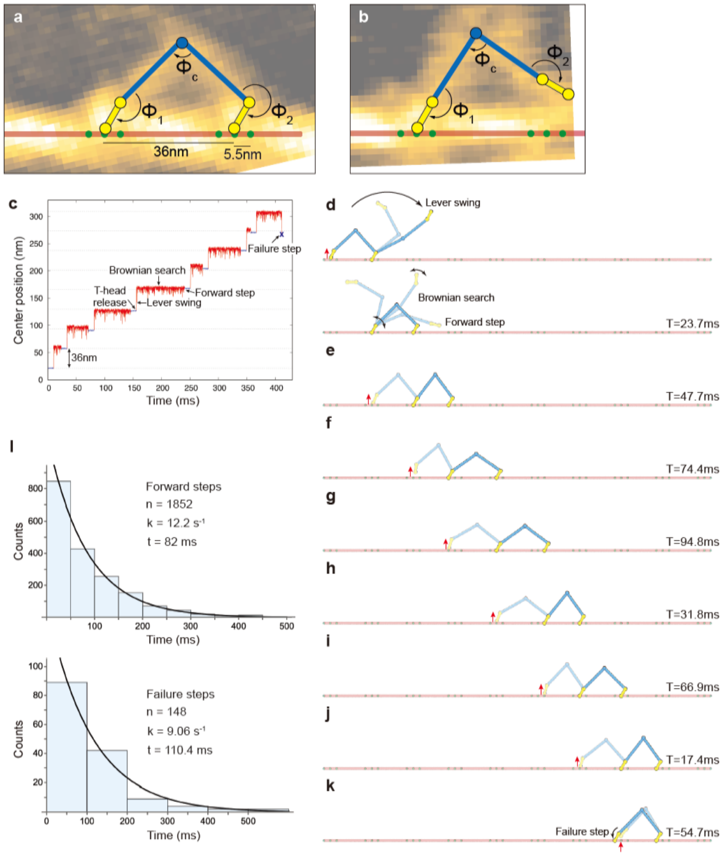


**Supplementary Fig. 10 | Mechanical model and simulations of ATP-free walking along actin filaments. a,b** Mechanical model of the M5 dimer in the double-headed (**a**) and single-headed (**b**) bound states. Motor and neck domains are represented as rods connected by beads, characterizing an instantaneous conformation by the indicated three angles $\phi_1$, $\phi_2$ and $\phi_c$. HS-AFM snapshots in the double-headed and single-headed bound states are shown as references in transparent. Numerical simulations were performed for the dynamics of the indicated 5 beads. The actin filament was viewed as a rod with available M5 motor domain binding sites (green circles) separated by a step size of 36 nm or 36 ± 5.5 nm. **c** Simulation of ATP-free M5 walking. Shown is the time trace of the dimer (central bead position) for eight consecutive steps. After a waiting time of 10 ms (blue lines), each step proceeded by detaching the T-head from its actin binding site followed by lever swing and Brownian search (red lines) until rebinding to actin occurred. The eighth event was a failure step. **d–k** Corresponding conformational snapshots of the M5 model. T-head detachment is indicated by the red arrows. In (**d**), the lever swing and Brownian search processes are highlighted by several snapshots in transparent. For each step, panels (**e–k**) display the M5 conformation shortly after T-head release (in transparent) and after complete rebinding to actin (in opaque) (see Supplementary Video 6). Landing times for each step are given. **l** Statistics of landing times. The histograms show the distribution of landing times for forward step (top) and backward return bottom), acquired from a simulation of 2,000 M5 steps. The rates obtained from exponential fitting (black lines) are given.

**Supplementary Methods**

**Modelling of myosin V dimer.**

In the HS-AFM experiments, motions of the entire M5 dimer are practically confined to proceed over the planar lipid bilayer surface. To model the lever-arm swing and Brownian search in an ATP-free translocation cycle, one needs to formulate a stochastic dynamical 2D description for M5. To obtain estimates for the stepping time in the absence of ATP from numerical simulations, a simplified model would suffice.

Our analysis of the HS-AFM video images does not reveal any significant stretching or bending of the M5 necks. Therefore, in contrast to other models[1,2], we approximate that they are rigid and remain straight. Since the motor domains may also be treated as rigid bodies, an instantaneous configuration of a dimer can be characterized by two neck–motor domain hinge angles and the central neck junction angle (**Supplementary Fig. 9a**). Moreover, we assume that when a motor domain is bound to actin, it maintains a fixed orientation with respect to the actin filament. While one might also treat necks and motor domains as physical rods, an even simpler description was used in our model for the M5 dimer to determine viscous friction forces acting on it. Namely, we view the dimer as consisting of five beads that are connected by very thin, straight and stiff, links. Only the beads, not the links, experience viscous friction when they move through the fluid; the mobilities of all beads are taken as the same. This simplification allowed us to perform Brownian dynamics simulations of a five-bead M5 model.

**Modelling of actin filaments.**

Within the simplified modelling scheme, the actin filament is described as a static rigid rod. Along it, binding centres for M5 motor domains are placed periodically with a spacing of 36 nm. In the helical shape of actin filaments, several neighbouring binding sites are accessible to motor domains. To roughly take this into account in our two-dimensional model, we introduced to each binding centre a left and a right neighbours, each separated by 5.5 nm from the central spot. Thus, each interaction site available for a motor domain corresponds to a triple of point-like binding centres (**Supplementyary Fig. 9a**).

**Modelling of interactions between M5 and actin filaments.**

We have to account for three situations. The first is a motor domain bound to actin at a specific site. In our model, a bound motor domain is characterized by its corresponding two beads having pinned positions. The end bead has the same position as that of the binding centre, and the second bead has a fixed position such that the motor domain orientation is 60° with respect to the actin filament. The second situation corresponds to the detachment of a bound motor domain from the actin filament. This is modelled by unpinning the corresponding motor domain beads. Eventually (the third situation), an unbound motor domain should be able to rebind to one of the actin binding spots. It is assumed that binding of a motor domain to actin can occur when two conditions are satisfied (**Supplementary Fig. 9b**). First, the end-bead of a motor domain should be at a distance not more than $r$ from the nearest binding centre on the actin filament. Second, the relative motor domain orientation with respect to the filament should not deviate by more than the angle $\Delta$ from the fixed orientation of 60° of the bound motor domain. Once these conditions are reached, the binding event instantaneously take place. The choice of the interaction radius $r$ is related to the Debye length of electrostatic interactions, and that for the tolerance angle $\Delta$ consideres orientations that would correspond to rebinding to either site of the actin binding triple.

**Additional model constraints.**

(i) We prevented crossing of M5 with the actin filament. This was implemented by always requiring $Y_i(t) > 0$ for all beads; the actin rod had coordinates $Y = 0$.

ii) We also roughly prevented self-biting of the two heads by imposing a restriction on the central hinge angle $\phi_c$, requiring always $16° < \phi_c < (360° - 16°)$.
iii) For an unbound M5 head, we introduced a period $T_{\mathrm{delay}}$ during which rebinding to the same site from which it became unbound was prevented. We used $T_{\mathrm{delay}}$ = 35 ms. All model parameters and values used in the numerical simulations are provided in **Supplementary Table 1**.

**Simulation course of ATP-free M5 stepping by T-head unbinding.**
We considered two different sets of simulations: i) a single simulation demonstrating a few stepping cycles and ii) 2,000 independent single-step simulations to accumulate statistics of landing times. In (i), the initial condition for the short simulation was the 5-bead model with the T-head bound to actin at the centre of one binding triple and the L-head bound at the centre of the next binding triple. Unbinding of the T-head was initiated after a waiting time $T_{\mathrm{u}}$ = 10 ms, chosen for visualization purposes only. After rebinding of the unbound motor domain to either of the binding triples at the forward site or to one of the binding triples at the backward site (where the T-head became unbound), one step was finished. The next step was initiated, again after the waiting time $T_{\mathrm{u}}$, by unbinding the T-head. The simulation was terminated after 8 completed steps. In (ii), the initial condition for each single-step simulation was the 5-bead model with the T-head motor domain bound to actin at the centre of a binding triple, and the L-head motor domain bound at the centre of the next binding triple. Unbinding of the T-head was initiated at the start of the simulation at time $t$ = 0. After rebinding of the unbound motor domain to either spot of the binding triple at the forward site, or to one of the binding triples at the backward site (where the T-head became unbound), a single step was finished and the landing time was recorded. Thus, we have accumulated a distribution of step times.

**Supplementary Table 1. Parameters and their values used for mechanical model and simulations of ATP-free M5 translocation.**

| Parameter | Value | Notes |
|---|---|---|
| **Model parameters** | | |
| Motor domain length $l_h$ | 7 nm | Ref [3], SI text |
| Neck length $l_n$ | 25.5 nm | SI text |
| Converter hinge | | Fitting of HS-AFM data |
| Equilibrium neck–motor domain angle $\phi_0$ | 148.5° | |
| Stiffness $k_0$ | 23 $k_B T$ | |
| Stiffness $k_1$ | 2.3 $k_B T$ | |
| Parameter $b$ | 34.4° | |
| **Binding parameters** | | |
| Bound-motor domain orientation | 60° | Ref [3], SI text |
| Actin site separation $L$ | 36 nm | Ref [4], SI text |
| Separation between adjacent actin sites $L_t$ | 5.5 nm | Ref [5], SI text |
| Rebinding delay (perturbed site) | 35 ms | see SI text |
| Binding radius $r$ | 2.1 nm | see SI text |
| Binding tolerance angle $\Delta$ | 18° | see SI text |
| **Numerical parameters** | | see Methods/SI text |
| Rod stretching stiffness $\kappa_s$ | 150 | |

| | |
|---|---|
| Integration time-step | $5 \cdot 10^{-4}$ (1.4 ns) |
| Central hinge angle constraint | $16^\circ < \phi_c < 344^\circ$ |
